# Oligodendrocyte-mediated Myelin Plasticity and its role in Neural Synchronization

**DOI:** 10.1101/2022.07.03.498570

**Authors:** Sinisa Pajevic, Dietmar Plenz, Peter J. Basser, R. Douglas Fields

## Abstract

Temporal synchrony of signals arriving from different neurons or brain regions is essential for proper neural processing. Nevertheless, it is not well understood how such synchrony is achieved and maintained in a complex network of time-delayed neural interactions. Myelin plasticity has been suggested as an efficient mechanism for controlling timing in brain communications through adaptive changes of axonal conduction velocity and consequently conduction time delays, or latencies; however, local rules and feedback mechanisms that oligodendrocytes (OL) use to achieve synchronization are not known. We propose a mathematical model of oligodendrocyte-mediated myelin plasticity (OMP) in which OL play an active role in providing such feedback. This is achieved without using arrival times at the synapse or modulatory signaling from astrocytes; instead, it relies on the presence of global and transient OL responses to local action potentials in the axons they myelinate. While inspired by OL morphology, we provide the theoretical underpinnings that motivated the model and explore its performance for a wide range of its parameters. Our results indicate that when the characteristic time of OL’s spike responses is between 10 and 40 ms and the firing rates in individual axons are relatively low (⪅ 10 Hz), the OMP model efficiently synchronizes correlated and time-locked signals while latencies in axons carrying independent signals are unaffected. This suggests a novel form of selective synchronization in the CNS in which oligodendrocytes play an active role by modulating the conduction delays of correlated spike trains as they traverse to their targets.

**Significance Statement:** Synchronization of signals arriving from different neurons or brain regions is of great importance for proper brain function. In vertebrates, myelination provides an efficient way to achieve this by adjusting the conduction velocity and consequently the conduction delays. It is increasingly evident that myelination is an adaptive process and involved in learning; however, the local rules and the feedback that guide oligodendrocytes towards achieving desired delays are not known. Here we propose a simple and biologically plausible model of myelin plasticity in which oligodendrocytes play an active role in providing the needed feedback. We use theoretical arguments, mathematical modeling and computer simulations to show its robust synchronization capabilities which enable efficient communication between the brain regions.

---

**T**emporal precision required in neural processing can range from sub-millisecond in sound localization and echolocation tasks to milliseconds and hundreds of milliseconds in perceptual and motor system signal processing. Often, this is a consequence of individual neural cells or brain regions requiring a narrow time window to integrate signals arriving from multiple sources. Signals traveling from distant regions commonly traverse complex conduction paths along which conduction velocity (CV) is not constant and undergoes dynamical changes and perturbations, particularly during development. This will alter arrival times of action potentials which may undermine the required temporal precision. It has been argued for more than a decade that a solution to this problem is the adaptive adjustment of conduction velocity through a mechanism of *myelin plasticity* (MP) (1, 2), which postulates that myelination is an adaptive and neural activity-dependent process. Modifying myelin sheath thickness and node of Ranvier structure provides the most efficient means to alter conduction delays through changes in conduction velocity. There is growing evidence that myelin plasticity is important for fear conditioning(3, 4), spatial learning(4, 5), and is shown to be essential for motor skill learning(6–9). Yet, very little progress has been made in understanding the local learning rules in this new form of plasticity and what feedback may be used by oligodendrocytes (OLs) to adjust myelination in the central nervous system (CNS). OLs are mostly located far from the target neurons and lack direct feedback on what the desired CV is, because the information about the spike arrival times is not available at these intermediate locations. In most studies of *activity-dependent* myelination (ADM) (10–14) the precise timing of individual spikes is ignored. It is well known that the introduction of time delays can change both stability and synchronizability at a system level, which then provides an indirect mechanism for MP to affect both the stability (10) and synchronization, for example in a network of FitzHugh-Nagumo neurons(14). These schemes are based on the local activity rate in the connections and do not explicitly include spike timing information in their local rules.

In principle, the arrival times at the target can be explicitly used as the feedback signal, via learning curves similar to that of spike-timing-dependent plasticity (STDP)(15) but with some important differences. In STDP, the crucial parameter is the pre- and post-synaptic spike time difference, Δ*t*, the sign of which determines whether long-term potentiation or depression occurs, with Δ*t* = 0 marking the sharp transition between the two. For MP, such learning curves would be unstable and hence must be ramped across the Δ*t* = 0 line (16, 17). More importantly any such feedback information at the target will have to be passed in a retrograde fashion, which is problematic since OLs are mostly located very far from the post-synaptic targets of the axons they myelinate. The same problem applies to schemes in which a network of oscillators (Kuramoto) is studied and the feedback is based on phase differences(18).

To develop spike-timing-dependent myelination (STDM) rules, it becomes important to consider schemes in which the mediators of feedback have to act locally and base adjusting the delays only on local signal timing information, where the information about the arrival time error is not available. In this work, we propose models in which OLs are not only the myelinating agents but also serve as the mediators, providing feedback through the interaction with spikes in different axons. We call this form of STDM *oligodendrocyte-mediated myelin plasticity* (OMP). Specifically, we focus on a particular type of OMP, which uses the spike-response of oligodendrocytes and its transient temporal profiles as a reference for adjusting CV and relative time delays. We use theoretical arguments, mathematical modeling and simulations to show that even the simplest form of OMP models (OMP-1) can lead to effective synchronization of correlated and time-locked signals, while leaving temporally uncorrelated signals unaffected.

## Oligodendrocyte-mediated Myelin Plasticity (OMP)

Our OMP model was inspired by OL morphology, which differs drastically from that of Schwann cells (SC), which myelinate axons in the peripheral nervous system (PNS). The most important morphological difference between these two myelinating cell types is that the OL extends many of its processes to myelinate multiple axons, while the SC myelinates only a single axon (fig. 1A,B). The OL connectivity is schematically depicted in fig. 1C and quantified via the myelination matrix, 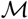 (see Methods). A single OL can have up to 50 such processes extended to axons within a 100-200 *μm* distance from the soma, but with a tendency to maximize the number of axons it can myelinate, making it unlikely that a given OL would myelinate the same axon twice(20, 21). We postulate that this morphology enables a single OL to integrate signals from different axons and act as a mediator in providing the needed feedback for adjusting the relative timing of different signals via dynamic regulation of the CV along these axons. In fig. 1D we outline a general form for such spike-response OMP models, consisting of three essential steps, described below. In these continuous-time models, the transient nature of the temporal profile of the OL response to an action potential plays a crucial role in creating the needed reference for adjusting the relative delays on different axons.

**Fig. 1.**
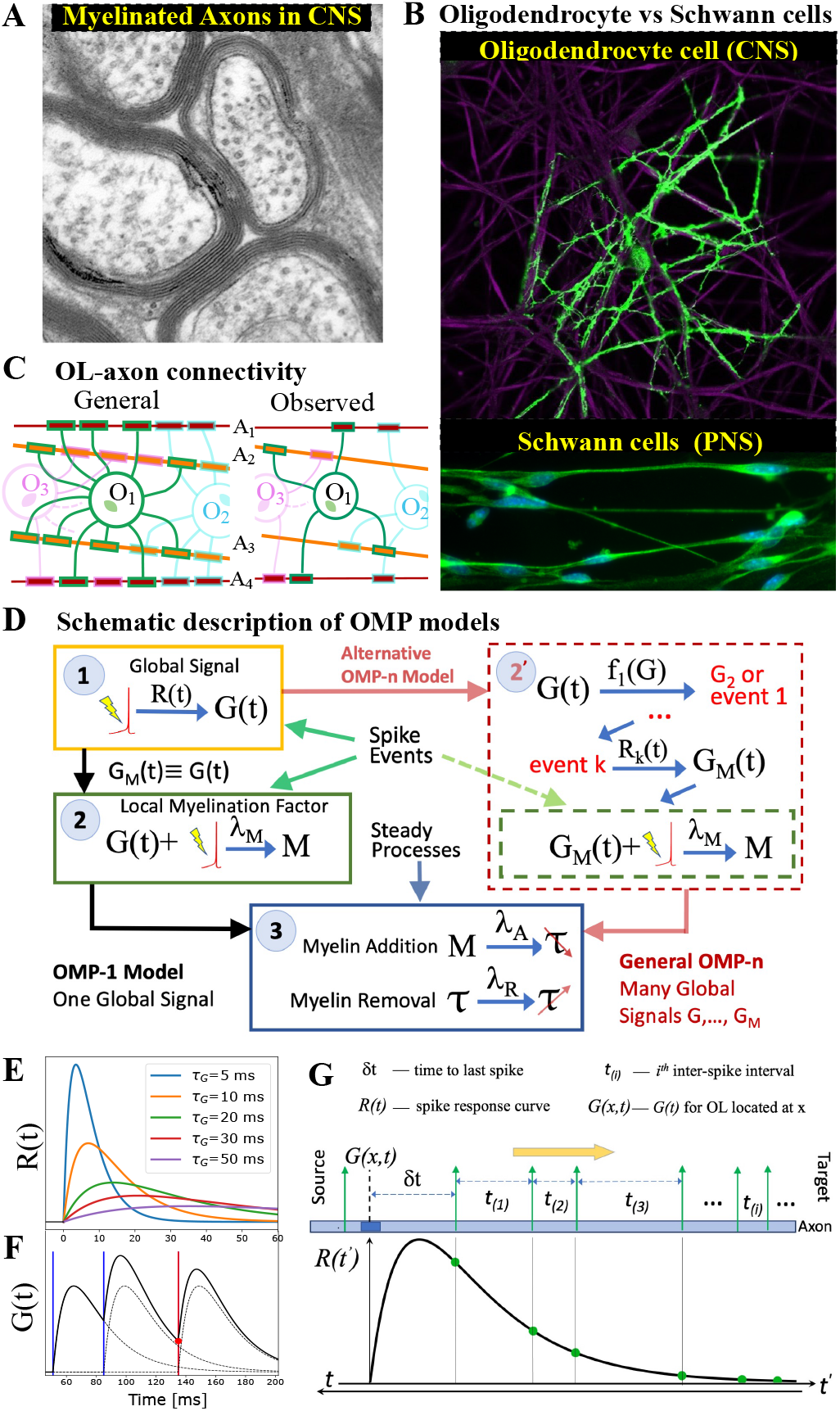
OMP motivation and model description. **(A)** Cross-section of myelinated axon bundle in CNS illustrates that myelin thickness on adjacent axons differs; (B) single OL (green) myelinates many different axons (purple), which is in stark contrast to Schwann cells in the PNS, which myelinate only one axon; (C) OL-axon connectivity: OL tend to avoid placing multiple processes on a single axon, thus maximizing the number of axons it myelinates; (D) schematic depiction of OMP models. In general, they can have a complex cascade of global signals and events, each with its own response function, *R_k_*(*t*) (dashed red box). Here, we study OMP-1 which has only the three basic steps required for OMP models to work; (E) spike-response curves *R*(*t*) for increasing values of *τ_G_*; (F) the equation governing the release of *G*(*t*) is linear and the response to multiple spikes (vertical lines), is the linear sum of individual responses (dashed lines). The release of *M* after each spike at any given OL process/axon will be proportional to the amplitude of *G*(*t*) at the time of spike (red dot for red spike); **(G)** the sum over responses can also be viewed as sampling of *R*(*t*) (in reverse time) which we use in our theoretical derivations. A and B reprinted with permission from (19).

The first essential element of OMP is the release of a *global* intracellular signaling factor, *G*, after each neural spike on any of the axons myelinated by a given OL. This response has a characteristic transient temporal profile, *R*(*t*) (Step 1 in fig. 1D). When the OL encounters a sequence of spike trains, the resulting global intracellular signal, *G*(*t*), will fluctuate in time and provide a common and time-dependent reference to all of its processes. To allow for differential myelination between different axons, a *local* myelin promoting factor, *M*, is also required for each of the OL processes, which selectively modifies the CV of that axon. The *M* is also released after each neuronal spike (Step 2), but its release is catalyzed by *G* and hence is proportional to the global signal, *G*(*t*). In more elaborate OMP models (red dashed box, Step 2′ in fig. 1D), the release of *M* can depend on a different global factor, *G_M_*, which potentially is released via a cascade of events triggered by the original factor *G*. Here, we use the simplest model of this kind, OMP-1, which has only one global signal, *G*, that does both, responding to neuronal spikes and modulating the local release of the myelin-promoting factor, *M_a_*, at axon *a*, i.e., *G_M_* ≡ *G*. The last step (Step 3) represents two continuous processes, one being the conversion of *M* into myelin with some rate, λ_*A*_, (myelin addition) and the second being the steady removal of myelin with rate λ_*R*_ (12).

### OL equation and spike-response curves

The spike response curve, *R*(*t*), represents the global response of an OL to a single neuronal spike on any of its myelinated axons, which gives OMP its timing-dependent character. In a more general OMP model there can be several such curves responding to any triggering event in a cascade of responses, as depicted in fig. 1D. Each of the responses represents the change in a concentration or amplitude of some global factor released in the OL. For *R*(*t*) we chose a form that is an impulse response of a linear system, as described in the Methods. This form can have different rise and decay characteristic times, *τ_r_* and *τ_d_*, but here we use a single characteristic time of the OL spike response, *τ_G_* = *τ_r_* = *τ_d_*, yielding a simple form for *R*(*t*),

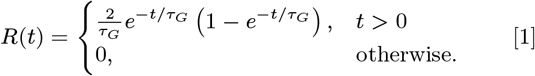

In fig. 1E we show a set of such curves for varying values of *τ_G_*. The equation for *G*(*t*) that gives such an impulse response is,

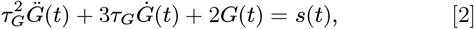

which governs the global response of an OL to input *s*(*t*).

OMP-1 is a continuous time model and the input signal, *s*(*t*), appearing in eq. (2), can technically be any integrable input. However, the time-dependent MP will have to rely on events that are sharply defined in time. We use trains of neuronal spikes for individual axons which are prescribed using a particular interspike interval (ISI) distribution, *p*_ISI_(*t*_(*i*)_), i.e., they are generated with a renewal process. An important parameter that characterizes these trains is their mean firing rate, *f_s_*, or equivalently the mean ISI, *τ_s_* = 〈*t_i_*〉_*t*_ = 1/*f_s_*. We use two main forms for *p*_ISI_ distributions: (a) the exponential distribution with the rate *f_s_* = 1/*τ_s_* with added fixed refractory time, *t_R_*, yielding the refractory Poisson process and (b) regular spiking, spaced at constant intervals, *τ_s_*. These *p*_ISI_s cover two extremes: for Poisson spiking, with *t_R_* = 0, the appearance of the next spike is completely independent of the previous spikes (memoryless process), and for regular spiking, the appearance of the next spike is precisely determined by the last spike. To both of the forms of *p*_ISI_ we also add *jitter*, specified by *σ_j_*, which spreads the spike times, such that these temporal shifts are normally distributed according to 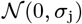. Such a mix of refractory Poisson and regular spiking with added jitter seems to cover the spike dynamics for communication between many areas of the brain (22).

When the spikes on different axons follow the same renewal process, i.e., obey the same *p*_ISI_ distribution, we call these “pure” signals, and among them distinguish two different cases: (1) the renewal processes for different axons are fully independent, and (2) they are time-locked via imposed relative time shifts between spikes in different axons, that we call *fixed delays*, 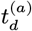, *a* = 1,…, *N_A_* (see section C in SI). The fixed delays represent either consistent temporal delays of signals coming from different sources or result from structural perturbations in the conduction pathways. Due to added jitter, which is always independent between different axons, the spikes will not be precisely time-locked, but will still be correlated. In fig. 2A, we show examples of correlated and independent spike trains, prior to the imposition of fixed delays, for both, Poisson and regular spiking.

**Fig. 2.**
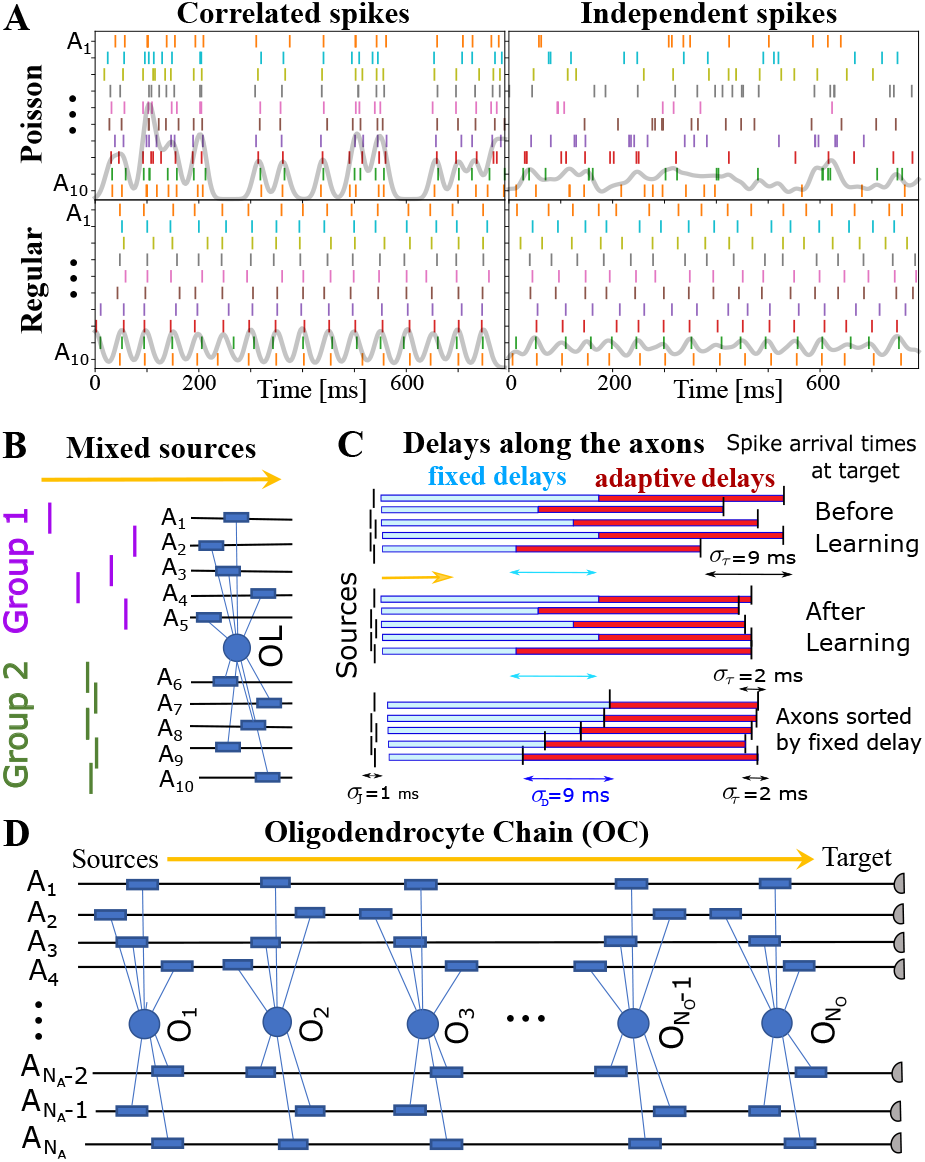
OMP simulations with spiking signals conducted along oligo-chain. (A) examples of “pure” spiking signals used in our simulations: correlated *versus* independent signals, and Poisson versus regular spiking. Gray lines are a Gaussian 10 ms window averages of all spikes; (B) example of “mixed” signals in which Group 1 of axons is conducting independent spikes and Group 2 time-locked/”correlated” spikes. The two groups can potentially interfere with the expected behavior of each “pure” group, i.e., affecting OMP’s ability to synchronize the “correlated” group of signals, or erroneously synchronizing the independent signals. Different groups could also contain spikes time-locked within each group, but independent between the groups; (C) schematic depiction of time delays between the signal source and the target. Fixed delays are not modifiable and essentially represent the spread in spike times as they enter the axonal bundle of myelinated axons, whose delays are adaptive. Note, that the horizontal bars represent the magnitude of the fixed and adaptive delays, and not axons; (D) OC with *N_O_* “effective oligodendrocytes” (segments), myelinating *N_A_* axons (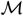 is an *N_O_* × *N_A_* matrix of ones).

The spike trains are generally formulated as a sequence of Dirac delta functions, 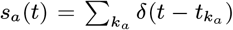, where *t_k_a__* indicates the time of the *k^th^* spike on axon *a*, and the sum goes over all spikes. Then, the solutions of the linear system in eq. (2) can be obtained analytically and the resulting *G*(*t*) can be written as,

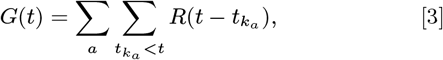

which is simply the sum of the responses to spikes on all axons that it myelinates, where the sum goes only over the spikes that happened prior to a given time *t* (fig. 1F). In our work we do not use eq. (3) directly, but rather solve eq. (2) numerically. The details of our simulation and implementation are described in section C.

### OMP-1 Model Learning Equations

The fluctuating global signal, *G*(*t*), obtained via eq. (2), serves as a catalyst for the local myelination promoter, *M*. We model this by making the increase in *M* proportional to *G*(*t*), as well as to the signal strength in the axon it myelinates. Since the myelination promoter, *M*, is also continuously converted to myelin with some rate, λ_*A*_, the differential equation for *M* on axon *a*, *M_a_*, can be written as,

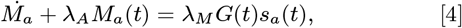

where λ_*M*_ specifies the production rate of *M* (fig. 1D,table S1) and it is the same for all processes. It is an OMP parameter that effectively controls the rate of adaptive changes in our model. The presence of the promoter in a given axon will lead to the increase in myelin sheath thickness (with rate λ_*A*_), and will compete with another continuous process of myelin removal which, in the absence of any activity, decreases with some rate, λ_*R*_.

For most neuronal plasticity models, saturation functions need to be introduced to stabilize the learning process. Similarly, in our implementation of OMP, we introduced two separate saturation functions for myelin addition, 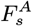, and removal, 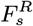. The OMP equation for the latency on axon *a*, *τ_a_*, can be written as

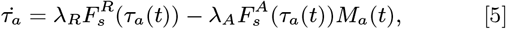

where 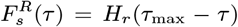 and 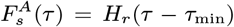, *H_r_*(*x*) = *xH*(*x*)/(*τ*_max_ – *τ*_min_) is the normalized ramp function, *H*(*x*) is the Heaviside (unit step) function, *τ*_max_ and *τ*_min_ are the parameters of the model which specify the maximal and minimal delays that are attainable on any axonal connection, respectively.

In Methods, we describe modifications to this basic model, one being the case of λ_*A*_ → ∞ (*instantaneous myelination*), for which eq. (4) is not needed, and another case includes a homeostatic equation (eq. (15)), which presumes that overall myelination for each OL reflects a long-term homeostatic steady-state between adding and removing myelin.

### OMP Theory

While OL morphology inspired our OMP model, the motivation arises from basic theoretical considerations that also provide hints about model performance. For uncorrelated signals, due to symmetry, any of the axons is equally likely to produce a spike, resulting in equidistributed concentrations of *M* guided only by the average concentration, *G*_av_ = 〈*G*(*t*)〉_*t*_. Predicting the synchronization effects for time-locked signals is a more difficult task, due to presence of jitter and non-linear saturation effects, particularly when the OL myelinates many axons. Nevertheless, we can make some simple predictions for the concentration of *M_a_* for Poisson spiking and for *N_A_* = 2 with regular spiking.

For individual axons, we presume the inter-spike intervals (ISI) are independent and identically distributed (i.i.d.) with density, *p*_ISI_(*t*_(*i*)_) (renewal process) and the resulting *G_a_*(*t*) can be estimated by recognizing that the sum over spikes in eq. (3) can equivalently be seen as sampling the *R*(*t*) in reverse time (see fig. 1G). Here, the important parameter is the time to the last spike, *δt*, as the remaining spikes are just the samples drawn from *p*_ISI_(*t*_(*i*)_), effectively making *G_a_*(*t*) a function of *δt_a_* = min |*t* – *t_k_a__*|, ∀*t_k_a__* < *t*. If we label the cumulative sum of the subsequent interspike intervals as 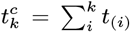, the expression for *G_a_*(*t*) can be written as

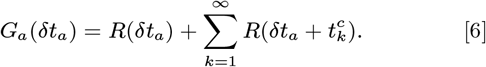

When combining *G_a_* from all axons, mutual ordering of spikes will generally need to be considered. For example, for the case of two axons (*N_A_* = 2) and a given fixed delay, *t_d_*, we need to consider separately the cases where *δt* ≤ *t_b_* = *τ_s_* – *t_d_* and where *δt* > *t_b_* (see fig. S1A), in order to derive the expressions for *G*_av_, 〈*M*_1_〉, 〈*M*_2_〉, as a function of *τ_s_*, *τ_G_*, and *t_d_*. For *τ_s_* < 2*t_d_*, the “leading edge” axon becomes the follower, in which case the myelination pattern is reversed, making its CV faster instead of slower (fig. S1B and C). The calculations become more cumbersome with increasing *N_A_*, even when jitter is ignored, which is true for most of the renewal processes governing the spiking dynamics on a given axon. However, in the case of a pure Poisson process (*p*_ISI_(*t*) = *e*^−*t*/*τ_s_*^, *t_R_* = 0), we can utilize its memoryless property to derive simple expressions for *G*_av_ and 〈*M_a_*〉. The expression for *G*_av_, at any of the locations along the axons is simply

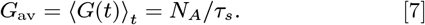

The expectation for the myelin promoter for the pure Poisson spiking (*t_R_* = 0) is also simple. When *N_A_* = 2, the average myelin promoter concentration at the leading edge axon is 〈*M*_1_(*t*)〉 = *C_M_G_av_* while for the lagging axon 〈*M*_2_(*t*)〉 = *C_M_*(*G_av_* + *R*(*t_d_*)), where *C_M_* = λ_*M*_/(λ_*A*_*τ_s_*). Hence, the ratio of their concentrations is always lower than one,

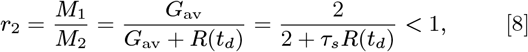

which will consistently lead to a greater increase in CV of the lagging axon, hence supporting synchronization. This also reveals the geometry of OMP synchronization, expected to be most efficient for *r_2_* ≪ 1, which happens at low firing rates (large *τ_s_*) and a fast spike response time, *τ_G_* (fig. S1C). In the case of multiple axons, the above argument retains its simplicity and we can write the expected concentration of the promoter *M* on axon *a* as

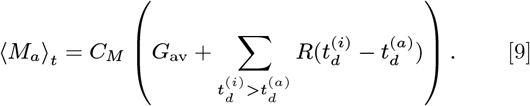

The equation 9 is an important result which indicates that with fixed time delays along different axons, the differential expression of the myelination factor will always myelinate the lagging axons more than the preceding ones. The expression in eq. (9) assumes no jitter, but still agrees well with the simulated values for *σ_j_* < 3 ms (fig. S2).

### OL Chain

The consistency of eq. (9) can, for small jitter and fixed delays, cause instabilities as the “leading” axon will keep losing myelin more than other axons. Generally, the spikes on less delayed axons will produce less *M_a_* than spikes on more delayed ones, creating an imbalance. This trend will stop only after the rate of change is significantly slowed down by the saturation limits and the decreasing λ_*R*_, due to homeostatic regulation. This problem is not surprising as most of the plasticity models are inherently unstable (e.g., Hebbian) and rely on saturation mechanisms. In the case of OMP models, this problem can also be resolved by using a natural assumption that the final arrival time will depend on the action of all OLs along a given axon bundle; the temporal differences of spikes across the bundle will become smaller for OLs close to the target compared to OLs close to the sources. For this reason it is worth, if not necessary, to simulate the sequential action of multiple OLs, in which the preceding OL-axon bundle segment can pass its modified spike arrival times to the next segment. Hence, we simulate a sequence of OMP equations, each feeding its output to the next segment. We call this an *oligo-chain* (OC), which is graphically depicted in fig. 2D. The OC will have *N_O_* sets of OMP-1 equations, each having their own myelin promoters and their own delays. As already emphasized, the OC depicted in fig. 2D does not imply literally that there are *N_O_* OL cells along the axons, but rather that there are *N_O_* segments, representing *N_O_* different populations of oligodendrocyte cells myelinating different portions of the axonal bundle, which modulate the delays locally. Assuming that all cells within the same segment will receive the same pattern of spikes and respond to it in the same way, they all can be governed by a single OMP equation. Individual oligodendrocyte cells, in fact, would not be able to modify the delays effectively and independently from other oligodendrocyte cells in the same location, as they would not be able to form tight nodes of Ranvier, considering that OLs prefer not to myelinate the same axon multiple times. Neighboring OL cells are then needed to stack their processes, reducing the width of the nodes of Ranvier and in this way greatly increasing the conduction velocity, i.e., reducing the conduction delays. Hence, we have a sequence/chain of *N_O_* “effective” OL cells, each modifying its local fraction of the total delay along *a*^th^ axon, 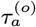, so that a total conduction delay on the axon is just the sum of all local delays 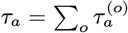.

## Results

We show in fig. 3A the estimated distribution of the long-time baseline parameter, 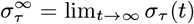, as well as the model selection chart, when fitting a large number (n=17280) of synchronization profiles, *σ_τ_*(*t*), obtained using a wide range of OMP parameters (see table S2a,figs. S3 and S4 for details). The results indicate that the OMP model behaves as desired. When the myelinated axons conducted correlated spikes, we observed a significant reduction in the conduction delay spread, *σ_τ_*, i.e., a significant increase in synchronization. No change was observed for independent spikes, and synchronization profiles were best fit to the constant model, C (purple) (section D, fig. S4). For correlated spikes, a single exponential approach to synchrony was most commonly observed (E1). In several instances, *σ_τ_*(*t*), was not monotonic and sometimes appeared oscillatory, which can be seen upon inspecting individual runs, particularly for small *N_O_* (*N_O_* < 3), short *τ_G_*, and small jitter, *σ_j_* (fig. 3B). As suspected, the OMP model with only a single OL has an inherent instability due to the presence of fixed delays, which could be alleviated by increasing *τ_G_* and *σ_j_*. For longer chains, however, this instability rapidly disappeared for any level of jitter (fig. 3B, solid lines; see also figs. S8 and S10). These trends can be seen in fig. 3C, where the averages of all *σ_τ_*(*t*) profiles are shown (table S2b, grouped by *N_O_*). The runs for small *N_O_* were longer, to match them in terms of the number of spikes processed. When plotted in actual time, it becomes evident that having more OLs in an OC greatly increases OC stability as well as the synchronization/learning rate, *L_τ_*. Since the averages were obtained over a large number of runs, the standard error (SE) is small and the oscillatory or other forms of instabilities average out. To better quantify all *σ_τ_*(*t*)s we fit them to five different models, as described in Methods and section D. In fig. 3D we show the 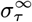 distributions and the model selection chart for different values of *N_O_*. They all indicate that, with increasing *N_O_*, synchrony is improved and instabilities disappear. For *N_O_* = 10 most of the OMP simulations yielded a stable exponential decay to a synchronized state (73% of all runs reduced the arrival time spread from the initial 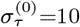 ms to below 3 ms; for *τ_G_*=10 ms this increases to 97%, with 74% synchronized below 1 ms).

**Fig. 3.**
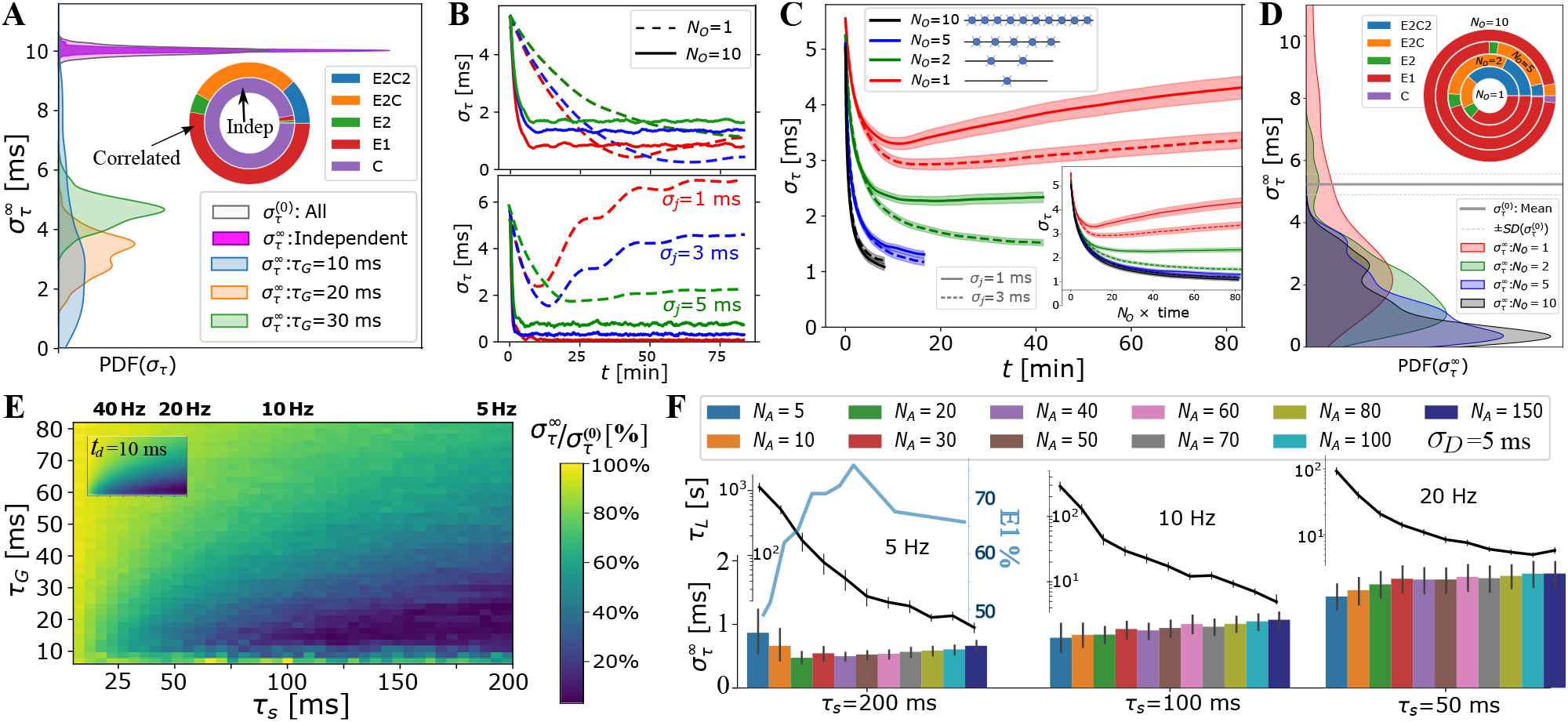
OMP-1 model behavior as a function of OMP and signal parameters. (A) When signals are independent *σ_τ_*(*t*) consistently show no change in overall synchronization (purple; innermost circle in the model selection pie-chart); for correlated signals significant synchronization occurs, largely depending on *τ_s_* and *τ_G_*;(B) individual OMP synchronization profiles, *σ_τ_*(*t*), simulations for *N_A_* = 10, λ_*H*_ = 10^−6^, λ_*M*_ =0.02, with *τ_G_*=30 ms (top panel) vs. *τ_G_*=10 ms (bottom). Dashed lines are for *N_O_* =1 and solid lines for *N_O_* = 10. Colors indicate the jitter level: *σ_j_* = 1 ms (red), *σ_j_* =3 ms (blue), *σ_j_* =5 ms (green). (C) dependence of *σ_τ_*(*t*) on the number of OLs, *N_O_*, in OC. We show both the comparison based on the raw time, as well as when matched in terms of the total number of neural spikes encountered by the OLs (inset); (D) density of 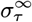 estimates dependence of *σ_τ_* on the number of OL in OC; (E) Percent reduction in *σ_τ_* vs *τ_G_* and *τ_s_* for *σ_D_*=10 ms; (inset) ratio of the promoter expression for two axon case, *N_A_* = 2, which matches well the overall pattern of synchronization;(F) dependence of 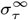 (bar plot) and the synchronization time constant, *τ_L_* = 1/*L_τ_*, (semi-log plot, above) on the number of axons, *N_A_*. *τ_L_* values are only for runs declared as E1 (blue curve inset indicates percent of E1, for different *N_A_* (for all spike rates). OMP parameter values explored in this figure are listed in table S2

For correlated spikes, the effectiveness of synchronization strongly depended on two temporal parameters: the characteristic response time, *τ_G_*, and the mean inter-spike interval, *τ_s_*, which was assumed to be the same for all axons. In fig. 3E, we explored the synchronization effect as a function of *τ_G_* and *τ_s_*; these runs used *N_O_*=5, *σ_j_*=1, see table S2c. For very short *τ_G_* < 10 ms, performance was unstable, reflecting the fact that for the short-lasting spike responses, *R*(*t*), the comparison window between spikes in different axons is too narrow. On the other hand, having spike responses last too long, i.e., *τ_G_* > 40 ms, makes the resulting *G*(*t*) too smooth to differentially release *M*, particularly for small *τ_s_* (S1C)). Accordingly, we found that for intermediate values of *τ_G_* ∈ [10 – 40] ms, synchronization was predictably achieved and was highly efficient for firing rates *f_s_* < 10 Hz.

In fig. 3F, we study the dependence of 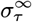 and the synchronization time constant, *τ_L_* = 1/*L_τ_*, on the number of axons that OLs myelinate, *N_A_*. The results are grouped by the firing rate simulated. For a low firing rate of 5 Hz, optimal performance was achieved for *N_A_* = 20, whereas synchronization became progressively harder with increase in firing rate and number of axons to synchronize. In general, increasing *N_A_* sped up synchronization for all firing rates explored. The fastest synchronization achieved at 5 Hz for an intermediate number of axons (30 < *N_A_* < 60) also coincided with the highest fraction of stable, E1, synchronization profiles (peaking at *N_A_* = 50, light-blue). We note that myelinating more than *N_A_* = 50 axons did not improve synchronization efficiency for any of the spiking rates, which, perhaps, hints to why OLs rarely extend more than 50 processes.

In fig. 4A-C, we tested the ability of the OMP model to selectively handle mixed sources of signals by having non-overlapping groups of axons carry spikes with different *p*_ISI_, or different mutual correlations. Spikes from different groups are inducing responses in the same OL, and hence mutually interfere and can potentially corrupt the expected behavior for the equivalent “pure” group. In fig. 4A, B, we show the results when one group of axons carries correlated and another carries independent spikes. In fig. 4C, we explore the effect of different groups carrying signals that are correlated within each group, but not between groups. We evaluate *σ_τ_* within each group and compare it with the equivalent pure signal sources, the number of axons being matched in all comparisons. We found an increase in synchronization only for correlated groups, with only slight “jamming” of the synchronization performance due to presence of other groups with potentially corrupting signals. The independent signals remain unaffected, which is a desired behavior for selective synchronization of spikes from different neuronal populations.

**Fig. 4.**
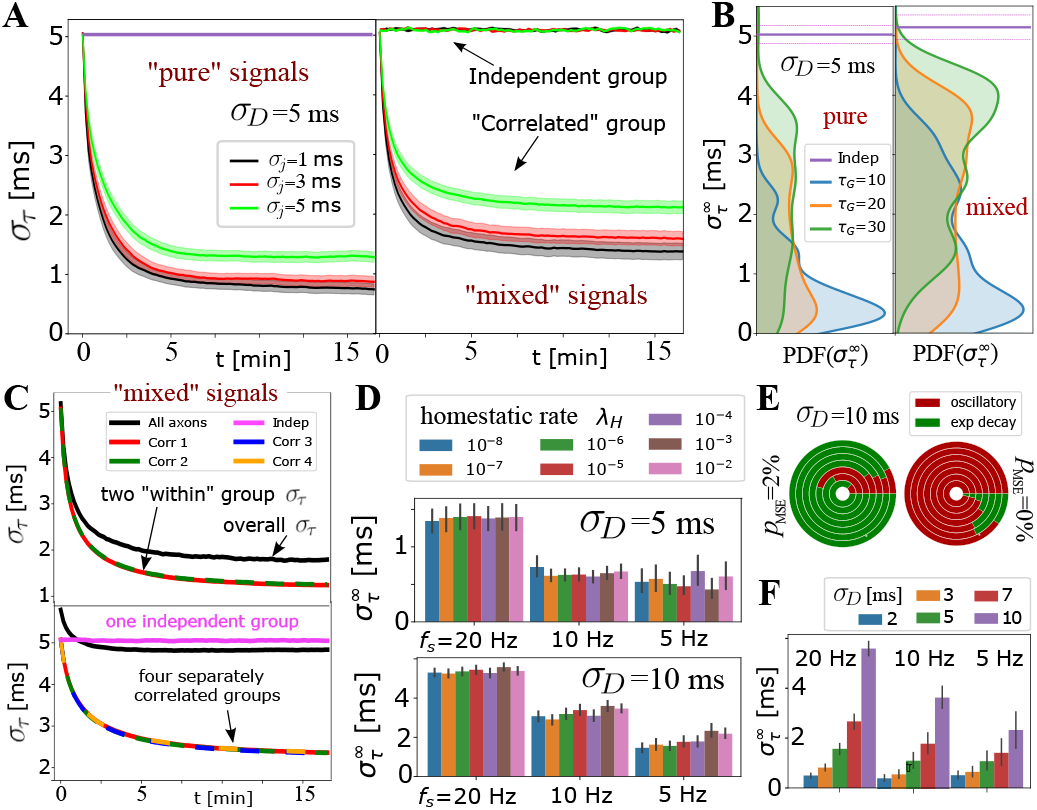
Selective synchronization with mixed signals; effects of λ_*H*_ and *σ_D_* parameters. (A) Average *σ_τ_*(*t*) (grouped by *σ_j_*; shaded regions indicate SE) over many parameter values for *N_A_* = 10, *σ_D_* =5 ms. In the left panel is the average for the set of *N_A_* = 10 axons conducting only “pure” correlated signals, and on the right are the averages within two groups of 10 axons, out of *N_A_* =20 total that OL myelinates, with “mixed” signals – one group conducting. The behavior of the correlated groups is clearly distinguishable from the independent ones (indicated by arrows); (B) estimated density for 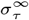 comparing pure (left) vs mixed signals (right panel) for a wider range of parameters, including *N_A_* =20 and *N_A_* =50, but with number of axons carrying correlated signals matched in all cases (*N_A_* = 10, *N_A_* = 25). (C) top panel shows *σ_τ_*(*t*) for two groups of correlated signals (colored lines) with Poisson spiking (correlated within group, but mutually independent); bottom panel shows results for 4 separately correlated groups + 1 independent group (cyan). Black lines represent the spread over all axons; (D) investigating the influence of homeostatic regulation and homeostatic rate, λ_*H*_, grouped by different firing rates and different *σ_D_*; (E) model selection chart depicting the relative proportions of oscillatory (E2C2, E2C) vs stable, single exponential (E1) *σ_τ_*(*t*), for two different values of the F-test fudge-factor *p*_MSE_; the innermost circle is for λ_*H*_ = 10^−2^ and the outermost for λ_*H*_ = 10^−8^; (F) OMP synchronization performance for different magnitudes of fixed delays, *σ_D_*. OMP parameter values used for the simulations in this figure are listed in table S3.

For OMP to be operational requires an overall balance between myelin removal (controlled by λ_*R*_) and myelin addition (controlled by λ_*M*_ and λ_*A*_). For example, if λ_*R*_ is too large the long-term behavior of the OMP model would lead to complete myelin removal. Here we achieve such an operational regime by treating λ_*R*_ as a variable and control its rate of change with the homeostatic rate, λ_*H*_, as defined in the homeostatic equation (eq. (15)). The results in fig. S6C indicate that this homeostatic regulation, while keeping the model operational, does not play an essential role in OMP synchronization. Although it appears that for low firing rate and for large fixed delays (*σ_D_* = 10 ms), increasing the value of λ_*H*_ has some detrimental effect on model performance, this effect is not seen for *σ_D_* = 5ms. The oscillatory dynamics of λ_*R*_ depends on λ_*H*_ and λ_*M*_ (see fig. S10) which can propagate into synchronization profiles. Nevertheless, the pie-chart in fig. 4E does not indicate that λ_*H*_ strongly influences the model selection, independent of what value of *p*_MSE_ was used. Figures S8 and S10 show that the oscillations roughly average to the same value and thus do not significantly affect synchronizability in long term. We note that the average values of λ_*R*_ deviate from the expected value 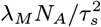 even for the case of independent spikes (fig. S9). This difference arises from ignoring saturation effects in eq. (14) and depends on the values of *τ*_nom_, *τ*_min_, and *τ*_max_. We also explored scenarios in which we use the “true” balancing homeostatic value of λ_*R*_, obtained via a trial run, and set λ_*H*_ = 0. The corresponding synchronization profiles were not significantly affected, further indicating that the homeostatic process is not an essential element for achieving spike synchronization in the OMP model.

The production rate of *M*, λ_*M*_, while greatly influencing the learning rate, *L_τ_*, has negligible influence on 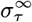 (fig. S6A,B). Similarly, the conversion rate λ_*A*_ had very little influence on the outcome of OMP (fig. S6D), indicating that the simplified OMP-1 model with instantaneous myelination could be a more efficient way in studying its behavior (fig. S3E). In fig. 4F, we explored the dependence of 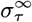 on the magnitude of fixed delays, *σ_D_*. For correlated signals, synchronization always occurs but its efficiency decreases in terms of 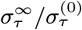 when *σ_D_* becomes large (for *σ_D_* > 10 ms see fig. S5F). Such large delays might be commensurate with the delay corrections needed during the development, but are presumably much larger than the timing corrections needed in the adult brain.

## Discussion

Here we report the development of a simple, biologically plausible model of oligodendrocyte-mediated myelin plasticity that synchronizes temporally correlated neuronal spikes as they travel along an axonal bundle. This enables the OLs in the model to counteract the temporal dispersion arising from heterogeneous conduction delays and allow synchronous arrival times of spikes coming from distant neuronal populations with correlated activity. While the general idea of myelin plasticity is not new(1, 2), a local mechanism by which the brain could robustly and selectively adjust axonal latencies has been missing. Our OMP model introduces robust local learning rules and feedback mechanisms for adaptive changes that yield desired results under a wide range of biologically realistic parameters.

The adaptive changes to axonal delays are selectively applied to groups of axons that carry correlated spikes so that they arrive at their targets simultaneously, while those carrying independent or uncorrelated spikes are not affected (fig. 4A-C). This selectivity conforms to known relationship between circuit anatomy and function, in which neighboring axons share similar temporal firing and functional properties; for example, the tonotopic organization of auditory cortex and the cortical homunculus in somatosensory cortex. Such correlated firing also drives refinement of connections between the retina and LGN during development(23). Myelination usually begins after axons reach their target and become functional, starting from the cell body and proceeding toward the axon terminal. This is clearly evident in the optic nerve, where OL progenitor cells migrate out of the brain and into the optic nerve during development, yet axonal myelination proceeds in the reverse direction, beginning at the retina on retinal ganglion cells and proceeding toward the optic chiasm (24). Such a proximal-to-distal gradient agrees well with our OMP model as the OLs at the source end of the OC will experience less synchronized signals. Previous work that use phase- and time-dependent models of myelin plasticity(17, 18) presume *a priori* that the temporal difference feedback at the target is available to OLs, however, this local information at the target, i.e., synapse, will have to be transported in a retrograde fashion. Besides being slow, such process would suggest that more myelin will be found close to the target rather than far away from it. We note, that with the existence of targeted fast axonal transport, e.g. via mitochondria, it is still possible that a promoter is transported in a retrograxde fashion far from the synapse.

The fact that OLs with more than 50 processes are uncommon is also in accord with our observation that having *N_A_* > 50 is not advantageous, according to our OMP results. Other testable prediction of this model would be that structural deformation or lesion to a portion of axons carrying correlated activity would lead to re-myelination downstream or immediately after the site of lesion, while myelin upstream will not be affected. Results in fig. 4 suggest that the efficiency of synchronization is slightly reduced when a fraction of axons carry uncorrelated signals. Hence, we postulate that in the brain OLs might be removing their processes from such axons, while keeping or growing new processes only on those axons that carry synchronized signals. OMP-1 predicts that synchronization is most effective at lower firing rates, and thus we expect it to occur when the brain operates in low spiking rate regimes.

It was observed that OLs can undergo sudden depolarization which leads to significant changes (10%) in conduction velocity (25), which might suggest that such events are essential for efficient MP. While this can be incorporated into a general OMP scheme (fig. 1D), the OMP-1 shows that passive responses of OL cells are sufficient for synchronizing neural signals, as well that astrocytes, commonly considered the actors in providing the feedback in MP, are not needed. Our results also demonstrate that the presence of noise acts as a stabilizer, both in terms of jitter removing the instabilities in the OMP model as well as Poisson spike dynamics always biasing larger concentrations of the *M* to be on the axons with larger fixed delays, as opposed to regular spiking which can reverse this pattern and destabilize the system. Hence, Poisson spiking, even though slower and less effective in many situations than the regular spiking, is a more reliable form of MP. Our model of synchronization provides further support for the narrative that the noisy brain is a healthy brain, and too much regularity/synchronization in the brain dynamics can lead to its failure.

The OMP-1 model described here, while simple, is not the simplest form that can serve as a proof of concept. An instantaneous myelination model described in Methods, also shows robust synchronization performance (fig. S3E), with shortcomings coming only at high learning rates (large λ_*M*_) and when using homeostatic regulation in eq. (15), making it then a stochastic equation. But, we kept a more general form, since we expect that in the future developments a more sophisticated regulation of λ_*M*_, λ_*A*_ and λ_*R*_ will be needed, to account for elaborate time-locking patterns where the firing frequencies are different but matched via integer multiples. One of the weaknesses of the current model is that λ_*M*_ is constant and the same for all axons, making the model sensitive to rate differences, which can override the synchronization effects. To make it work, the production rates, λ_*M*_, need to have their own homeostatic regulation, so that the axons with higher firing rate will down-regulate their λ_*M*_. In a more elaborate OMP models a mix of activity- and time-dependent MP might be needed. The net result will depend on the relative strengths/learning rates between the time-dependent and the activity-dependent learning, something that will require its own independent study. For the current model, we explored its sensitivity to firing rate variability (fig. S6E,F) which showed that variations in firing rate greater than 5% are highly detrimental to OMP’s ability to adjust fixed delays properly. When the firing rates across axons are very different, it might also not be desirable to synchronize those groups of axons, nor is easy to define the temporal synchrony in such situation.

There are other aspects of OMP that are not explored here. For example, the effects of inhomogeneous OL-axon connectivity is not addressed, as we use the same myelination matrix along the OC. The stochasticity in our models comes mainly from the stochasticity of the spikes and their jitter. Future work will address complex patterns of connectivity, other independent sources of noise for both global and local factors, as well as more sophisticated homeostatic regulation discussed above. Delays in factor *G* are mainly implemented here through the rise time, *τ_r_* = *τ_G_*, however, increase in *G* coming from a particular OL process will not affect all processes simultaneously, and the relative delays between different processes can have significant effects on the resulting synchronization, which will need to be explored via delay-differential equations, or using a discrete implementation. Some of the proposed mechanisms for myelin plasticity are discrete in nature, for example the treadmilling model (26), but the discreteness is only in the state of myelination; the feedback mechanisms needed for temporal adjustments in any time-dependent MP might still need the continuous-time comparisons. Discretizing time in the simulations, can introduce its own effects, which for a long-time simulations required by the MP mechanism (MP time scale is many orders of magnitude longer than the neuronal spiking time scale) could dominate synchronization performance.

While the goal of OMP model was to provide a general mechanism by which synchronization can be achieved, our usage of the terms implies that *M* is a molecular factor, while *G* is some fast propagating global signal, e.g., intracellular potential or ionic concentration. Our model results suggest that the release and clearance of a global intracellular signal cannot be too slow since for *τ_G_* > 80 ms effective synchronization does not occur even for very slow firing rates. But it also suggests that *τ_G_*, the characteristic time for the release and clearance of *G*, does not have to be too rapid, and can even be detrimental in some situations. For firing rates below 10 Hz, *τ_G_* should ideally be in the range 10 ms < *τ_G_* < 40 ms for effective synchronization of correlated inputs even for large fixed delays (e.g., *σ_D_* = 10 ms). These timing requirements make intracellular Ca^2+^ a good candidate for the role of *G*, since it is also a catalyst for many bio-molecular reactions. Because of its diverse roles, it will require further investigation to suggest the identity of *M*(11, 27–29), which is outside the scope of the present work.

In summary, we demonstrated that a simple and biologically plausible adaptive dynamics of the OMP model leads to efficient and selective synchronization of correlated and time-locked signals, without affecting mutually independent streams. This is a novel perspective in brain organization in which white matter conduction properties and myelin plasticity act as a temporal “lens” tp “focus” multiple spike trains as they target a particular brain region. Our model also addresses lingering questions about the illusive local learning rules and feedback mechanisms in MP, as it circumvents the need for the direct information about the actual arrival times at the target. With its precisely spelled-out dynamics and learning rules it serves as a useful basis for designing future experimental tests aimed to elucidate the nature of MP in the CNS.

## Materials and Methods

### Spike-response curves

A general spike response curve is parameterized by separate rise and decay times, *τ_r_* and *τ_d_*; for a spike occurring at time *t_s_*, it can be written as,

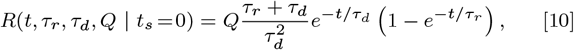

where *Q*, represents the single release amount/quantity, which is constant and independent of the current value of a global signal, *G*(*t*). The peak of the response is happening at time 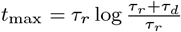, reaching the value 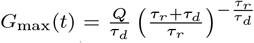. This form allows more general explorations (e.g., *τ_r_* ≈ 0, yields the exponentially decaying *R*(*t*)), but most importantly it is the impulse response curve of the second-order linear system,

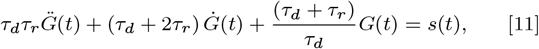

where *s*(*t*) represents the signal that drives the system, and *Ġ*(*t*) and 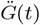 are the first and second time derivatives of *G*(*t*).

### OL-axon Connectivity

In general, each OL myelinates many axons and can have multiple processes on a single axon (fig. 1C), or can have none on others. But, the OL-axon connectivity can be described with the *myelination matrix*, 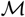, with OLs as rows and axons as columns, and indicates the number of processes a given OL places on each of the axons, e.g., for the “general” connectivity in fig. 1C, the matrix is,

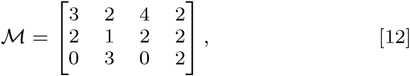

while the observed avoidance of OLs myelinating single axons with multiple processes(20, 21), makes 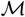 likely to consist of ones and zeros. Here, we use an “effective” OL, which represent a population of identical OLs and hence the issue of precise *OL-axon connectivity* is not as important. In a more elaborate model in which individual OLs are simulated, the issue of connectivity becomes more important. Such models will be computationally demanding, because of the combined effects of multiple OLs myelinating the same location along an axon. Since here we use effective OLs, we address only a simple situation in which a chain of *N_O_* OLs are fully myelinating *N_A_* axons. In this case 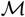 is simply an *N_O_* × *N_A_* matrix of ones. We assume stable connectivity and that modifications of CV are achieved only through remodeling of myelin sheaths and the nodes of Ranvier.

### “Instantaneous” Myelination

The conversion rate, λ_*A*_, appears not to play an important role (fig. S6D), particularly if OLs operate far from the saturation limits, as *M* is then just a currency for conversion into myelin, or, changes in CV. We can eliminate eq. (4) by taking the limit λ_*A*_ → ∞, i.e., presume that *M* is instantly converted to myelin, resulting in an immediate change in the time delay, *τ_a_*. The solution for *M_a_*(*t*) is obtained by convolving the expression on the right side of eq. (4) with the impulse response of the left side, *e^−λ_A_t^*. Using the fact that the signal on axon *a*, *s_a_* is a spike train, and replacing such *M_a_*(*t*) in eq. (5), we obtain

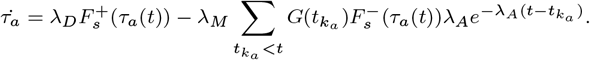

Since λ_*A*_*e*^−λ_A_(*t*–*t_k_a__*)^ can be interpreted as a Dirac delta function when λ_*A*_ → ∞, the “instantaneous” equivalent of eq. (5) becomes

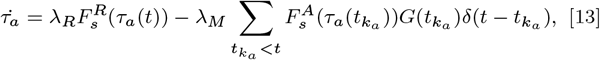

where it is indicated that *τ_a_*(*t*) in the sum will only depend on its values at spike times *t_k_a__*, after 13 is integrated. To have stable integration in this case, it is important to set λ_*M*_ sufficiently small.

### Homeostatic Regulation

The form in eq. (13) illustrates that the essence of the adaptive process for timing adjustments is the balance between continuous myelin removal (longer delay) and the discrete increments induced by spikes. For independent Poisson spikes and ignoring saturation effects, the balance condition is,

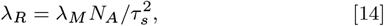

where *τ_s_* is the average inter-spike interval of the Poisson process on a single axon. However, with the saturation functions and when correlated signals are introduced, this homeostatic balance can be disturbed. In order to keep the system in balance, we make the removal rate of myelin, λ_*R*_, another time dependent variable in the dynamics of the OMP model. To do so we assume that each OL operates with some local and nominal homeostatic level of myelination, parameterized by some nominal delay, *τ*_nom_, such that any deviation from it will lead to a slow change in λ_*R*_ according to,

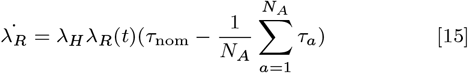

where λ_*H*_ is a homeostatic rate. We set it to a very small value (λ_*H*_ ≤ 10^−5^) so that the time scale for changes in λ_*R*_ is much slower than the time scale of individual spikes or the changes in myelination. In some instances, in order to test the importance of having eq. (15), we simply set λ_*H*_ to 0, after guessing or finding the value for λ_*R*_ that balances the increase in myelin content for a given input signal. This homeostatic rule can be interpreted as a tendency of each OL to conserve its overall amount of myelin, while re-distributing it over different axons. We assume that such an activity-dependent MP process is in place and simulate only its ability to adjust the overall rate of myelin removal, i.e. the myelin removal rate, λ_*R*_.

### OMP Simulations

We mainly focus on the ability of our OMP model to synchronize spikes with fixed temporal delays that arise either from developmental and other structural disturbances in the conduction pathways, or due to dynamically fixed temporal delays between different sources. The main measure we use is the final spread in the spike arrival times, as measured by *σ_τ_* = SD_*a*_ (*σ_D_* + *τ_a_*) (for details see section C). The spread starts with some large value, determined by fixed delays, 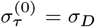, and in the course of time, due to the OMP mechanisms, is reduced to lower values, ideally to zero, indicating perfect synchronization. We analyzed its time-course during learning, *σ_τ_*(*t*) by fitting it to five nested models of the form,

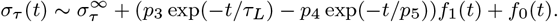

The two most basic models are constant(C), containing only the long-time baseline parameter, 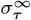, and the single exponential model (E1), which adds two more parameters, including a measure of the synchronization rate, *τ_L_* = 1/*L_τ_*. The double-exponential (E2) and two other models (E2C,E2C2), aiming to fit damped oscillatory behavior of *σ_τ_*(*t*) via functions *f*_0_ and *f*_1_ (with long-time limit 0 and 1, respectively), are used just to quantify the instabilities in learning. We introduce a fudge/tolerance parameter, *p*_MSE_, to modify F-test based nested model selection, and use 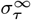, and/or *τ_L_* of the winning model to quantify the synchronization performance (section D). In the results, we plot the average *σ_τ_*(*t*) over a wide range of parameters, usually grouped by a given parameter of interest. Full details of our implementation and simulations are given in SI, including the glossary table of all parameters (table S1).

## ACKNOWLEDGMENTS

This research was supported by the Division of the Intramural Research Programs (DIRP) of the National Institute of Mental Health (NIMH), USA, (S.P., D.P., ZIAMH002797) and the Eunice Kennedy Shriver National Institute of Child Health and Human Development (NICHD), USA, (R.D.F.,ZIAHD000713; P.J.B, 1ZIAHD008972-04). This work utilized the computational resources of the NIH HPC Biowulf cluster (http://hpc.nih.gov).

## References

1. R Fields, White matter in learning, cognition and psychiatric disorders. Trends Neurosci. 31, 361–370 (2008).

2. RD Fields, Myelination: an overlooked mechanism of synaptic plasticity? The Neurosci. 11, 528–531 (2005).

3. S Pan, SR Mayoral, HS Choi, JR Chan, MA Kheirbek, Preservation of a remote fear memory requires new myelin formation. Nat. neuroscience 23, 487–499 (2020).

4. PE Steadman, et al., Disruption of oligodendrogenesis impairs memory consolidation in adult mice. Neuron 105, 150–164 (2020).

5. F Wang, et al., Myelin degeneration and diminished myelin renewal contribute to age-related deficits in memory. Nat. neuroscience 23, 481–486 (2020).

6. CM Bacmeister, et al., Motor learning promotes remyelination via new and surviving oligodendrocytes. Nat. neuroscience 23, 819–831 (2020).

7. D Kato, et al., Motor learning requires myelination to reduce asynchrony and spontaneity in neural activity. Glia 68, 193–210 (2020).

8. IA McKenzie, et al., Motor skill learning requires active central myelination. Science 346, 318–322 (2014).

9. L Xiao, et al., Rapid production of new oligodendrocytes is required in the earliest stages of motor-skill learning. Nat. neuroscience 19, 1210–1217 (2016).

10. S Pajevic, PJ Basser, RD Fields, Role of myelin plasticity in oscillations and synchrony of neuronal activity. Neuroscience 276, 135–147 (2014).

11. RD Fields, A new mechanism of nervous system plasticity: activity-dependent myelination. Nat. Rev. Neurosci. 16, 756–767 (2015).

12. DJ Dutta, et al., Regulation of myelin structure and conduction velocity by perinodal astrocytes. Proc. Natl.Acad. Sci. 115, 11832–11837 (2018).

13. B Stevens, S Porta, LL Haak, V Gallo, RD Fields, Adenosine: a neuron-glial transmitter promoting myelination in the cns in response to action potentials. Neuron 36, 855–868 (2002).

14. A Talidou, PW Frankland, D Mabbott, J Lefebvre, Learning to be on time: temporal coordination of neural dynamics by activity-dependent myelination. bioRxiv (2021).

15. Gq Bi, Mm Poo, Synaptic modifications in cultured hippocampal neurons: dependence on spike timing, synaptic strength, and postsynaptic cell type. J. neuroscience 18, 10464–10472 (1998).

16. CW Eurich, K Pawelzik, U Ernst, JD Cowan, JG Milton, Dynamics of self-organized delay adaptation. Phys. Rev. Lett. 82, 1594 (1999).

17. S Pajevic, PJ Basser, RD Fields, Models of plasticity and learning employing adaptive temporal delays in Annual Meeting of The Society for Neuroscience, Chicago, IL. (Soceity for Neuroscience), p. online (2015).

18. R Noori, et al., Activity-dependent myelination: A glial mechanism of oscillatory self-organization in large-scale brain networks. Proc. Natl. Acad. Sci. 117, 13227–13237 (2020).

19. R Fields, Regulation of myelination by functional activity. Neuroglia 3, 573–585 (2013).

20. L Dumas, et al., Multicolor analysis of oligodendrocyte morphology, interactions, and development with brainbow. Glia 63, 699–717 (2015).

21. DM Walsh, TD Merson, KA Landman, BD Hughes, Evidence for cooperative selection of axons for myelination by adjacent oligodendrocytes in the optic nerve. PloS one 11 (2016).

22. G Maimon, JA Assad, Beyond poisson: increased spike-time regularity across primate parietal cortex. Neuron 62, 426–440 (2009).

23. M Meister, RO Wong, DA Baylor, CJ Shatz, Synchronous bursts of action potentials in ganglion cells of the developing mammalian retina. Science 252, 939–943 (1991).

24. T Ishibashi, PR Lee, H Baba, RD Fields, Leukemia inhibitory factor regulates the timing of oligodendrocyte development and myelination in the postnatal optic nerve. J. neuroscience research 87, 3343–3355 (2009).

25. Y Yamazaki, et al., Modulatory effects of oligodendrocytes on the conduction velocity of action potentials along axons in the alveus of the rat hippocampal ca1 region. Neuron glia biology 3, 325–334 (2007).

26. DJ Dutta, et al., Regulation of myelin structure and conduction velocity by perinodal astrocytes (vol 115, pg 11832, 2018). Proc. Natl.Acad. Sci. 116, 12574–12574 (2019).

27. PM Paez, DA Lyons, Calcium signaling in the oligodendrocyte lineage: regulators and consequences. Annu. review neuroscience 43, 163–186 (2020).

28. H Wake, PR Lee, RD Fields, Control of local protein synthesis and initial events in myelination by action potentials. Science 333, 1647–1651 (2011).

29. H Wake, et al., Nonsynaptic junctions on myelinating glia promote preferential myelination of electrically active axons. Nat. communications 6, 1–9 (2015).

