## Supplemental Information for "Oligodendrocyte-mediated Myelin Plasticity and its role in Neural Synchronization"

#### This PDF file includes:

Supplementary text
Figs. S1 to S10
Tables S1 to S4
SI References

### Supporting Information Text

Our Supplementary Information (SI) contains supplementary text, followed by supplementary figures and supplementary tables of the parameter values used in our simulations. The Supporting Information Text portion is divided into five sections that are referred to throughout the main text with the following labels,

- OMP-1 Variables and Parameters ........ section A
- Implementation of OMP-1 ........ section B
- OMP-1 Simulations ........ section C
- Synchronization Profile Analysis: Model Fitting ........ section D
- Supplement to OMP theory ........ section E

**A. OMP-1: Variables and Parameters.** In its fullest form, the OMP-1 model can have up to 13 scalar parameters plus the $N_O \times N_A$ myelination matrix $\mathcal{M}$. We provide an overview of all OMP parameters in table S1, together with the OMP variables and other symbols used for specifying the spiking signals conducted along the axons and for the data analysis.

| List of symbols used in this manuscript | | |
|---|---|---|
| Symbol | Description | Value Range/Dimensionality |
| OMP-1 model parameters | | |
| $R(t)$ | OL response curve characterized by its rise ($\tau_r$) and decay ($\tau_d$) times | see $\tau_G$ |
| $\tau_G$ | single characteristic time for $R(t)$ ($\tau_G = \tau_r = \tau_d$) | [2-100] ms |
| $\lambda_M$ | myelin promoter production rate | 0.01 - 0.5 |
| $\lambda_A$ | myelin addition rate | 0.01 - 0.5 |
| $\lambda_R$ | myelin removal rate | $\approx \lambda_M N_A/\tau_s^2$; model variable when $\lambda_H > 0$ |
| $\lambda_H$ | homeostatic rate | [0, $10^{-5}$] |
| $N_O$ | number of OL in OC | [1-20] |
| $N_A$ | number of axons | [2-50] |
| $\mathcal{M}$ | myelination matrix | $N_O \times N_A$: {full connectivity, random} |
| $\tau_{\min}$ | minimal delay attainable on axons | 3 ms |
| $\tau_{\max}$ | maximal delay attainable on axons | 100 ms |
| $\tau_{\text{nom}}$ | nominal/homeostatic delay | [10-90] ms |
| $\tau_0$ | initial adaptive delays parameter | $\tau_0 = \tau_{\text{nom}}$ |
| $p_\tau$ | percent spread of initial time delays | 5% |
| OMP-1 model variables | | |
| $G(t)$ | global signal, one variable per OL (eq. (2)) | $N_O$ variables |
| $M_a(t)$ | local myelination promoting factor on axon $a$ (eq. (4)) | $N_O \times N_A$ variables |
| $\tau_a(t)$ | total time delay for axon $a$, or for particular OL ($\tau_a^o$) (eq. (5)) | $N_O \times N_A$ variables |
| $\lambda_R(t)$ | myelin removal rate as solution to eq. (14) | OMP parameter when $\lambda_H = 0$ |
| Spiking signal parameters | | |
| $\tau_s$ | inter-spike time interval parameter | [10-250] ms |
| $f_s$ | mean firing rate | [4-100] Hz |
| $t_R$ | refractory period for the case of refractory Poisson process | [0 - 100] ms |
| $\sigma_j$ | amount of jitter given to spikes | [0 - 10] ms |
| $\sigma_D$ | SD for fixed delays | [0 - 20 ] ms |
| $D_a$ | zero-mean fixed delays, value on axon $a$ | random values, $\mathcal{N}(0, \sigma_D)$ |
| $\sigma_s$ | percent variability in firing rates | [0 - 20] % |
| $T_{\text{exp}}$ | total duration of simulation | [10 min - 10 hrs ] |
| $n_e$ | number of epochs recorded during the whole simulation | [20 - 500 ] |
| $T_e$ | duration of epochs, after which parameters are recorded | $T_{\text{exp}}/n_e$ [10 sec - 5 min ] |
| $n_r$ | number of replicated simulations/trials | [3 - 10] |
| Quantification parameters/other symbols | | |
| $\sigma_\tau$ | OC spread SD($D_a + \tau_a$) | evaluated after each epoch; [ms] |
| $\sigma_\tau(t)$ | time-course of synchronization during OMP learning | $\sigma_\tau$ over all epochs; [ms] |
| $\sigma_\tau^{(0)}$ | initial spread before learning, $\sigma_\tau^{(0)} = \sigma_\tau(0)$ | determined by $\sigma_D$, $\tau_0$, and $p_\tau$ [ms] |
| $\sigma_\tau^\infty$ | long-time baseline, $\sigma_\tau^\infty = \lim_{t\to\infty} \sigma_\tau(t)$ | estimated via model fitting; [ms] |
| $\tau_L$ | characteristic time for synchronization | estimated via fitting; [min] or [s] |
| $L_\tau$ | learning/synchronization rate | $=1/\tau_L$ |
| $t_{(i)}$ | ISI for the $i^{th}$ interval | NA |
| $p_{\text{ISI}}(t_{(i)})$ | probability density function of ISI | NA |

**Table S1. List of symbols used in the manuscript: OMP model parameters, OMP variables, spiking signal parameters, and data analysis and quantification parameters and other symbols. For each we provide a short description and the range of the parameter values, or its dimensionality, in the case of the variables**

The nominal OMP-1 model is governed by three main equations: 2, 4, and 5, expanded with eq. (15), which adds a homeostatic control of $\lambda_R$, that now becomes an OMP-1 variable. Omitting the equation for homeostasis requires fine-tuning the balancing value for $\lambda_R$ for a given parameter setting. As shown in Methods, this model can be simplified further by assuming "instantaneous myelination", in which case eq. (4) can be omitted, when using eq. (13) instead of eq. (5); however, using eq. (13) requires, particularly if used together with eq. (15), that $\lambda_M$ be chosen sufficiently small to insure that the integration is stable. If $\lambda_M$ is large, sudden jumps in the value of $\tau_a$ will make eq. (15) stochastic. This can lead to negative values for $\lambda_R$ or even the delays themselves, which is not realistic.

The standard variables of the model, for a single OL, are $G(t)$, $M_a(t)$, and $\tau_a(t)$, $a = 1 \dots N_A$, giving a total of $2N_A + 1$ variables per OL, excluding $\lambda_R$. In practice, solving the model will require $2N_A + 2$ variables per OL, since eq. (2) is a $2^{nd}$-order differential equation, requiring an auxiliary variable (see section B). $G(t)$ and $M_a$ are "fast" variables that change on a milliseconds time scale (dictated by $\tau_G$ and impulse responses to the spikes, with mean ISI, $\tau_s$), while, $\lambda_R$ and $\tau_a, a = 1 \dots N_A$, are "slow" variables whose rate of change is controlled by $\lambda_H$ and $\lambda_A$, respectively ($\lambda_H \ll \lambda_A$). In fig. S7 we show an example of time progression for both fast (fig. S7A) and slow variables (fig. S7B), including the synchrony measure, $\sigma_\tau$ (top row) and the epoch-averages of $M_a$ (bottom row), when a set of correlated signals is conducted along OC. Examples shown are for an OL at the beginning of the oligo-chain with $N_O = 5$, $N_A = 10$, and and a single trial, except for $\sigma_\tau$ (dashed lines) and $\lambda_R$ (different colors), where the results from independent trials (3 total) are also shown. In fig. S10 we similarly show the time-progression for the unstable case, $N_O = 1$, as shown in fig. 3A ($\lambda_H = 10^{-6}$), but now for three different values of $\lambda_H$, indicating that the oscillations seen in $\sigma_\tau(t)$ are a result of homeostatic control. We note here, again, that $N_O = 1$ does not mean literally that there is a single oligodendrocyte acting on a given axonal bundle but rather a population of oligodendrocytes acting at a particular location along the axon receiving the same pattern of activations and responding to it in the same way. When independent signals are conducted along OC, the time progression of the slow OMP variables displayed only stochastic variations and no clear trends, as shown in fig. S9 (apart from initial "refocusing" of all $\tau_a$ to the same mean value, due to homeostatic constraints). This result was consistent across all simulations in which independent signals are used, with either Poisson or regular spiking $p_{\mathrm{ISI}}$.

**B. Implementation of the OMP-1 model.** The crucial element of our model implementation is solving a system of differential equations, 2, 4, and 5. To solve 2 we have in fact implemented a more general case, given by eq. (11), which can be written as a set of first order equations using an auxiliary variable, $v(t)$,

$$\dot{v}(t) = -a\,b\,G(t) - (a+b)v(t) + q_s\,\Sigma_{t_k}\delta(t - t_k), \qquad \text{[S1]}$$
$$\dot{G}(t) = v(t), \text{ with } G(0) = 0,$$

where constants, $a$, $b$, and $q_s$, are defined using the parameters of the general spike response curve (eq. (10), i.e., the characteristic rise, $\tau_r$, and decay, $\tau_d$, time constants, and the amplitude of the release, $Q$) as follows:

$$a = \frac{\tau_r + \tau_d}{\tau_r \tau_d},\; b = 1/\tau_d,\; q_s = \frac{Q(\tau_r + \tau_d)}{\tau_r \tau_d^2}.$$

To use 2 with single characteristic time, for simplicity, as is described in this manuscript, we set the characteristic rise and decay times to be equal, i.e., $\tau_r = \tau_d = \tau_G$ ($t_{\max} = \tau_G \ln 2$). We also set $Q$ =1 for all spikes, in which case the constants in eq. (S1) simplify to $a = 2/\tau_G, b = 1/\tau_G, q_s = 1/\tau_G^2$.

Implementation of eq. (4) is straightforward, as it just adds one more first order differential equation, while implementation of eq. (5) requires special handling of Dirac delta functions. The simplest way to do this is to integrate the OMP-1 equations between subsequent spikes, as the spikes are specified externally, and handle the discontinuities at each spike separately. For more sophisticated/alternative OMP schemes, which might require a cascade of events, as depicted in fig. 1D, some of which will require internally triggered events. In that case, everything needs to be implemented using some kind of "events" functionality (functions evaluated when a set of conditions on the variables are satisfied), which is available in many integration packages, as is the case with Python(1) solver in SciPy(2), *scipy.integrate.solve_ivp*, which is what we used. The full simulation code, including specification of parameters as well as subsequent data analysis, was implemented in Python. Depending on the parameters used, simulations took anywhere between a few hours to more than 10 days (including the repeats, $n_r \in [3, 10]$). Implementation of the same code in C, or using JIT (3), could make evaluations substantially faster, but might sacrifice some flexibility and ease in implementing the event handling.

**C. OMP-1 Simulations.** We conducted our simulations on a highly parallel National Institutes of Health Biowulf cluster (http://hpc.nih.gov) and divided them into more than a dozen studies. For each simulation study, we specify a set of values to be explored for each of the OMP parameters, as indicated in tables S2 and S3. In the early and exploratory phase, we coarsely identified the working range for the most important parameters and usually chose a few values, usually three, indicating the low, mid, and high values of its "working" range, and include sometimes explorations outside that range (as was done in our early, coarse exploration of the model). Due to a large number of parameters that could influence the performance, we could not afford to exhaustively explore all of them on a fine grid for a large range of values. Instead, in each study, we explored the influence of a particular parameter on a finer grid (see tables S2 and S3), or in some cases a group of parameters, for example, the exploration of $\tau_G$ and $\tau_s$, given in table S2c and shown in fig. 3E. Similar detailed explorations of other OMP properties were left for future work. For $\lambda_M$, we used four or five values covering two or more orders of magnitude.

In our simulations, we chose fixed delays to be randomly drawn from a normal distribution $\mathcal{N}(D_{\mathrm{mean}}, \sigma_D)$. Here $\sigma_D$ is the standard deviation among fixed delays $t_d^{(a)}$ on all axons and is an important parameter in our simulations, while $D_{\mathrm{mean}}$ is an arbitrary offset, insuring that the delays are positive. This positivity constraint is not very important since we were only interested in the spread of the synchronized spikes on different axons. These fixed delays are always combined with adaptive delays, i.e., the conduction delays, $\tau_a$, on myelinated axons, so we often normalize them to zero mean ($D_{\mathrm{mean}} = 0$), and we label such normalized fixed delay on axon $a$ as $D_a$. The spread of arrival times of synchronized spikes at the target will then simply be the standard deviation of $D_a + \tau_a$, i.e., $\sigma_\tau = \mathrm{SD}(D_a + \tau_a)$. We use $\sigma_\tau$ extensively in this manuscript, as a measure of such spread in arrival times, making it an inverse measure of synchronization with $\sigma_\tau = 0$ indicating perfect synchronization. In order to allow for an easier comparison between different sets of parameters, particularly when plotting the average $\sigma_\tau$ over many runs (see fig. S3A and the left panel in C), we introduce a normalization step, so that the initial spread due to fixed delays is exactly $\sigma_D$, i.e., $D_a^{\mathrm{norm}} = \sigma_D D_a / \mathrm{SD}(D_a)$.

We initialized a set of $N_A$ random fixed delays, parameterized by their spread, $\sigma_D$, for each trial ($n_r$ total). We initialized the two $N_O \times N_A$ matrices, one carrying the information about $M$ concentration for each OL process in OC, and the other carries the local delays on each axon for each OL in the OC. The former is initialized to zero (all $M_a = 0$) and the latter is initialized based on the specified mean, $\tau_0$ and the percent spread, $p_\tau$, i.e., $\tau_a \tilde{\sim} \mathcal{N}(\tau, p_\tau \tau/100)$. For simplicity, we set $\tau_0 = \tau^o_{\mathrm{nom}} = \tau_{\mathrm{nom}}/N_O$, and we used $p_\tau$=5%. We created $N_A$ spike trains with prescribed dynamics, based on a given $p_{\mathrm{ISI}}$, and parameterized by $\tau_s$ (examples shown, in fig. 2A). For time-locked signals, we generated a single train with a given $p_{\mathrm{ISI}}$, and others were shifted versions of the same, based on the values of fixed delays ($D_a$, or $t_d^{(a)}$), which are subsequently randomized using jitter. The jitter spreads the location of all spikes, such that these temporal deviations are normally distributed according to $\mathcal{N}(0, \sigma_{\mathrm{j}})$. For each trial, we ran $n_e$ epochs of learning, each with duration $T_e$, and collected the values of the "slow" variables, $\sigma_\tau$, $\lambda_R$, as well as the mean value during the epoch of $\tau_a$ and $M_a$, for every OL in the OC. Due to the large number of runs we conducted, only $\sigma_\tau$ information was saved for every run, while other variables were saved only if the analysis requires it. When the homeostatic equation was used ($\lambda_H > 0$), we allowed 1 or 2 extra ("warm up") epochs to run, during which modifications to $\tau_a$ were disabled, allowing $\lambda_R$ to be closer to its equilibrium value when we start to track the spread, $\sigma_\tau$. Due to oscillatory behavior of $\lambda_R$, in most cases, this was not very important to do and did not affect $\sigma_\tau(t)$. In most studies, we obtained a large number of synchronization profiles, $\sigma_\tau(t)$, one for each parameter setting/set, which were then analyzed using our model fitting (see section D) or plotted vs time, usually as averages over many runs, grouped by variable(s) of interest (see fig. S3C).

**D. Synchronization Profile Analysis: Model Fitting.** The averages over large number of runs can obscure numerous details in synchronization profiles, $\sigma_\tau(t)$, obtained from single runs, such as instabilities. Hence, it is useful to summarize and condense thousands of obtained profiles in an automated fashion and obtain distributions of critical parameters, most importantly the long-time spread, $\sigma_\tau^\infty$, and the "learning time", $\tau_L$, which is the inverse of the learning rate, $L_\tau$. To do this, we use a rich model with 12 parameters which was able to capture most of those profiles reasonably well. In the main text, we wrote the general form of the full (unrestricted) model as,

$$\sigma_\tau(t) \sim \sigma_\tau^\infty + (p_3 \exp(-t/\tau_L) - p_4 \exp(-t/p_5)) f_1(t) + f_0(t), \quad \text{[S2]}$$

but here we spell out the exact fitting formula, where we have used cosine functions to implement oscillatory functions $f_0(t)$ and $f_1(t)$, with additional parameters describing their amplitude, period, and phase, as well as their damping factor. Hence, the most general fitting model we use, containing 12 parameters, is a double exponential function containing both, the multiplicative oscillation and damped additive oscillation. We label the full model as E2C2 (containing two exponential and two cosine functions), and its formula is

$$\sigma_\tau(t) \sim \sigma_\tau^\infty + (p_3 \exp(-t/\tau_L) - p_4 \exp(-t/p_5))(1 + p_6 \cos(2\pi t/p_7 + p_8) + p_9 \exp(-t/p_{10}) \cos(2\pi t/p_{11} + p_{12}) \quad \text{[S3]}$$

We then explore a set of restricted models, all nested within the full model above, starting with the simplest, constant model (C), having only one free parameter, $\sigma_\tau^\infty$, which, consequently is also contained in all other models, with progressively more parameters. These are: single exponential model (E1), double-exponential (E2), double exponential with multiplicative oscillation (E2C). The full set of models that we fit is then,

- **C:** $\sigma_\tau(t) \sim \sigma_\tau^\infty$
- **E1:** $\sigma_\tau(t) \sim \sigma_\tau^\infty + p_3 \exp(-t/\tau_L)$
- **E2:** $\sigma_\tau(t) \sim \sigma_\tau^\infty + (p_3 \exp(-t/\tau_L) - p_4 \exp(-t/p_5))$
- **E2C:** $\sigma_\tau(t) \sim \sigma_\tau^\infty + (p_3 \exp(-t/\tau_L) - p_4 \exp(-t/p_5))(1 + p_6 \cos(2\pi t/p_7 + p_8)$
- **E2C2:** full, unrestricted model, see eq. (S3)

We constrained the parameters during the fitting as follows: $0 \le \sigma_\tau^\infty \le 2\sigma_\tau^{(0)}$, $T_{\mathrm{exp}}/1000 < \tau_L < 1000 * T_{\mathrm{exp}}$, $0 < p_3 < 2 * \sigma_\tau^{(0)}$, $0 < p_4 < 2 * \sigma_\tau^{(0)}$, $T_{\mathrm{exp}}/1000 < p_5 < 5 * T_{\mathrm{exp}}$, $0 < p_6 < \sigma_\tau^{(0)}/2$, $T_{\mathrm{exp}}/25 < p_7 < \infty$, $0 < p_8 < 2\pi$, We used these limits to have better stability and to avoid extreme outliers.* An example of such fits is shown in fig. S4A. In most cases, they give very

*since the total number of runs was close to $100k$, we did not inspect every single fit, but only a small fraction of all fitted parameters values were obtained at the boundary

similar estimates for the most important parameter in the model, $\sigma_\tau^\infty$ (fig. S4A, but in many cases the estimates can differ substantially, depending on what model is selected (see (fig. S4B). Our desire is to opt for a simpler model, when possible, as it often provides more reliable estimates of $\sigma_\tau^\infty$, and in the case of E1, of $\tau_L$. This allowed us to categorize better the behavior of OMP and particularly to identify single runs in which the approach to synchrony was truly unstable, or oscillatory, or if $\sigma_\tau$ was diverging away from synchrony. This categorization was implemented by our model selection procedure described below.

***Model selection.*** All of the simpler/nested models we call restricted since they are essentially equivalent to the unrestricted model with the coefficients of all extra explanatory variables being restricted to zero. Of course, the unrestricted model, having more parameters will then always be able to fit the data better or at least as well as the restricted model, in terms of the mean-squared error (MSE). The question is, whether this improvement is sufficiently large to warrant sacrificing the level of parsimony of the restricted model. One approach to this problem is to use an F-test, which compares two models, the unrestricted (U) vs a particular restricted version (R). Given the obtained fits to $n_{\text{pts}}$ points for both models and their corresponding residual sum of squares (RSS) one can calculate the F-statistic, $F$, given by

$$F_{\text{num}} = \frac{\text{RSS}_R - \text{RSS}_U}{n_U - n_R}$$

$$F_{\text{den}} = \frac{\text{RSS}_U}{n_{\text{pts}} - n_U}$$

$$F = \frac{F_{\text{num}}}{F_{\text{den}}}$$

where $\text{RSS}_m$ is the residual sum of squares of model $m$. This $F$ is $F$-distributed, with $(n_U - n_R,\, n_{\text{pts}} - n_U)$ degrees of freedom, which we can use for our statistical tests. The null hypothesis in these tests is that the unrestricted model does not provide a significantly better fit than the restricted model, hence rejecting it at a given significance level makes us opt for a more complicated model.

Using the conventional F-test for model selection did not work well for our purposes, since the undulations we observe are nearly always statistically significant even if the most stringent tests with extreme significance levels were used, e.g., $p < 10^{-15}$. This was not surprising, as the synchronization profiles were never truly statistically constant or exponential. For example in figure S4C, when $\sigma_\tau(t)$ is observed on a full scale, one would expect that the constant model is the best description of the behavior observed. However, the model selection with regular F-test chooses the less restricted models, in fact E2C2 in this case. After zooming in, we see that the results were indeed not constant, as those deviations from constancy were not just due to noise, but were significant. If the obtained simulation profiles were noisier then the restricted models would have a reasonable chance of being selected. Instead of adding artificial noise to our $\sigma_\tau(t)$, we introduced a parameter, $p_{\text{MSE}}$, with which we essentially control what level of noise or deviations we deem tolerable. The $p_{\text{MSE}}$ expresses this level of tolerance as the percentage of the initial, $\sigma_\tau$, or approximately, $\sigma_D$. Hence, we declare minimal amount of RSS in any fit,$\text{RSS}_{\text{min}} = n_{\text{pts}} * \text{MSE}_{\text{min}}$ and $\text{MSE}_{\text{min}} = (p_{\text{MSE}} * \sigma_\tau^{(0)}/100)^2$. This sets the level of MSE that is presumed by default, i.e., some minimal amount of noise present in residuals of any model. This is essentially specifying how much of RMS error can be tolerated in the restricted model, in order to rejected the null hypothesis that the unrestricted model is better. Since the numerator will remain unchanged, this essentially only modifies the denominator,

$$F^m_{\text{den}} = \frac{\text{RSS}_U + \text{RSS}_{\text{min}}}{n_{\text{pts}} - n_U},$$

where $\text{RSS}_{\text{min}} = n_{\text{pts}} * \text{MSE}_{\text{min}}$ and $\text{MSE}_{\text{min}} = (p_{\text{MSE}} * \sigma_\tau^{(0)}/100)^2$. The numerator portion is not affected, as it is a difference between two RSS. This yields the modified F-statistic, $F^m$,

$$F^m = \frac{F_{\text{num}}}{F^m_{\text{den}}} \qquad \text{[S4]}$$

that we use for our tests. This ad-hoc modification also changes the coefficients of determination for both models, $\text{R}^2_U$ and $\text{R}^2_R$, as follows,

$$\text{R}^2_U = 1 - \frac{\text{RSS}_U + \text{RSS}_{\text{min}}}{\text{TSS}_U + \text{RSS}_{\text{min}}}$$

$$\text{R}^2_R = 1 - \frac{\text{RSS}_R + \text{RSS}_{\text{min}}}{\text{TSS}_R + \text{RSS}_{\text{min}}},$$

if a different test, based on them is used. Here we use the modified F-statistic (eq. (S4), in most cases with $p_{\text{MSE}} = 2\%$. Under the modified test, the curve shown in fig. S4C is now declared as model C ("constant") for the three smallest significance levels that we use (see below).

Our $\sigma_\tau(t)$ consist of $n_{\text{pts}} = n_e$ data points which we use to estimate parameters for all models, via independent fits. We conducted a large number of randomly initialized fits, in order to insure that the best possible fit is obtained (fig. S4A and B). Note that due to the complexity of some of the models, the obtained fit of the unrestricted model, E2C2 might not end up finding the true global minimum and having the lowest MSE, but that happened very rarely (<0.3% or runs, and in all those cases E2C was the one with the minimal MSE). In our procedure, we start with the model with minimal MSE (usually E2C2), test it against all of the restricted models, and choose the most restricted model for which the null hypothesis is not rejected. We performed the tests separately at different but very low significance levels ($\alpha \in [0.01, 0.00001, 10^{-10}, 10^{-15}]$). The choice of the model was in many cases not strongly influenced by the choice of $\alpha$, but was strongly dependent on the choice $p_{\text{MSE}}$. For $p_{\text{MSE}} = 0\%$, even at extremely low $\alpha$, the unrestricted model would always be chosen (see fig. S4C). Using $p_{\text{MSE}} = 2\%$, allowed the choice of model to depend on $\alpha$ in most cases, as is indicated in fig. S4A, where E2C2, E2C, or E1 would be chosen, depending on how stringent the test was.

We chose $p_{\text{MSE}} = 2\%$, based on a set of 100 random examples, which we visually inspected and declared what the best model description is desired (e.g., the constant model in fig. S4C). Note that our model selection is largely ad-hoc and we emphasize that our modified F-test does not aim to provide a quantitative statistical analysis, as use of such absurdly small significance levels indicates, but only to provide a useful quantitative tool for summarizing tens of thousands of runs that we have performed. While the distribution of different models changes significantly for different choices of $p_{\text{MSE}}$ and $\alpha$, the derived values of the parameters $\sigma_\tau^\infty$ and $\tau_L$ is not significantly changed when different values of $p_{\text{MSE}}$ (but $> 1\%$) and $\alpha$ (but smaller than 0.01) were used. The same holds for the ad-hoc rule, of reverting to a less restricted model when $\text{MSE}_r > 500 \times \text{MSE}_U$, which happened very infrequently (see fig. S4D).

### E. Supplment to OMP theory.

***E.1. Derivation of instantaneous myelination. (iOMP-1) equation.*** We show here how eq. (4) can be reduced to eq. (13) by taking the limit $\lambda_A \to \infty$, i.e., by making the myelination instantaneous. In this way, we arrive at the simplest form of the OMP models, requiring only $3 + N_A$ parameters. Recapitulating equation 4 from the main text,

$$\dot{M}_a + \lambda_A M_a(t) = \lambda_M G(t) s_a(t), \tag{S5}$$

we have on the left a simple linear first order differential equation whose impulse response is $e^{-\lambda_A t}$, and hence the response to the input on the right can be obtained through the convolution,

$$M(t) = \lambda_M \int_0^t e^{-\lambda_A(t-t')} G(t') s(t') dt' + M(0) e^{-\lambda_A t} \tag{S6}$$

Replacing eq. (S6) in eq. (5), and using the fact that the signal on axon $a$, $s_a$ is a spike train we obtain

$$\dot{\tau_a} = \lambda_D F_s^+(\tau_a(t)) - \lambda_A \lambda_M F_s^-(\tau_a(t)) \sum_{t_{k_a} < t} G(t_{k_a}) e^{-\lambda_A(t - t_{k_a})} \tag{S7}$$

$$= \lambda_D F_s^+(\tau_a(t)) - \lambda_M \sum_{t_{k_a} < t} G(t_{k_a}) F_s^-(\tau_a(t)) \lambda_A e^{-\lambda_A(t - t_{k_a})}. \tag{S8}$$

We note that the expression $f_{\lambda_A}(t) = \lambda_A e^{-\lambda_A(t-t_{k_a})}$ can be interpreted as a Dirac delta function when $\lambda_A \to \infty$, since $\lim_{\lambda_A \to \infty} \int_{-\infty}^{\infty} g(t) f_{\lambda_A}(t-a) dt = g(a)$. The time dependence of $\tau_a(t)$ and $G(t)$ will then only depend on their values at spike times $t_{k_a}$, after integrating differential equation S8. Hence, the final form for this "instantaneous" equivalent of eq. (5) then becomes

$$\dot{\tau_a} = \lambda_D F_s^+(\tau_a(t)) - \lambda_M \sum_{t_{k_a} < t} F_s^-(\tau_a(t_{k_a})) G(t_{k_a}) \delta(t - t_{k_a}), \tag{S9}$$

which is also reported in the Methods and Materials portion of the main text (eq. (13)).

***E.2. Expectation values for $M_a$.*** Here we derive expressions for expected concentrations of the local factor, $\langle M_a \rangle$, in the case of no jitter, $\sigma_j = 0$. In the case of regular spiking, we do it only for $N_A = 2$.

***Poisson spiking.*** For a pure Poisson process, spike occurrences are independent from the time of the last or any previous spike, which is referred to as a memoryless property. If the rate of the Poisson process is $f_s = 1/\tau_s$, i.e., its ISI is governed by $p_{\text{ISI}}(t) = e^{-t/\tau_s}$, then the spikes are occurring within each infinitesimal time interval $(t, t + dt)$ with equal probability, $p_s(t) = dt/\tau_s$. Using this fact, we can integrate over all times in the past, and obtain the contribution from a single axon to be,

$$G_{\text{av}}(\delta t) = G_{\text{av}} = \int_0^\infty R(t) dt/\tau_s = Q/\tau_s. \tag{S10}$$

Because of its memoryless property, $G_{\text{av}}$ is independent of the temporal offset $\delta t$, and then the total contribution from $N_A$ axons is simply $G_{\text{av}} = N_A Q/\tau_s$. In the rest of the derivation (and the manuscript) we simply assume, $Q = 1$, without loss of generality, i.e., we show that $G_{\text{av}} = N_A/\tau_s$.

This is true for both cases, when the spikes are independent, or when they are arbitrarily time-shifted versions of each other, and we demonstrate the memoryless property here directly. In the case of independent Poisson processes this becomes trivial because the resulting spiking seen by the OL just becomes a new Poisson process with effective rate proportional to the number of axons $\tau_{N_A} = \tau_s/N_A$. In the case of correlated spikes, we demonstrate this by direct integration for the $N_A = 2$ case. We integrate first the period between 0 and $t_d$, when only the influence of the leading spike is seen,

$$I_1 = \int_0^{t_d} \frac{R(t')}{\tau_s} dt'. \qquad \text{[S11]}$$

After $t_d$, the "lagging" spike also releases $G$, and the expected value is

$$I_2 = \int_{t_d}^{\infty} \frac{R(t'-t_d) + R(t')}{\tau_s} dt'. \qquad \text{[S12]}$$

For our form for $R(t)$, these integrals evaluate to

$$I_1 = \frac{e^{-\frac{2t_d}{\tau_G}} \left(e^{\frac{t_d}{\tau_G}} - 1\right)^2}{\tau_s} \qquad \text{[S13]}$$

and

$$I_2 = \frac{2e^{-\frac{t_d}{\tau_G}} \left(\sinh\left(\frac{t_d}{\tau_G}\right) + 1\right)}{\tau_s}$$

Each integral individually looks cumbersome but the final result is the sum of the two, which becomes simply, $I = I_1 + I_2 = \frac{2Q}{\tau_s}$, which is what we wanted to show. This generalizes to all other delays yielding the expectation for $G_{\text{av}}$ to be $N_A Q/\tau_s$.

Next, we show that the expression for the concentration of the myelin promoter at any of the axons becomes also simple. We start with a burst of spikes with exponential ISI (Poisson bursts, with no refractory period, $t_R = 0$). For simplicity, we assume that there is no jitter, hence, the differences in timing are solely due to the fixed delays, $t_d^{(a)}$. At the occurrence of each spike, the myelin promoter is released proportional to the current value of G(t). For Poisson bursts, $G_{\text{av}}(t)$ is independent of $\delta t$, hence, the expected concentration of the myelin promoter for the leading edge axon (the least delayed axon) is simply $\langle m_{\text{fast}}(x)\rangle = C_M G_{av}(x)$, where $C_M = \lambda_M/(\lambda_A \tau_s)$. In essence, we have presumed here that the leading edge contains random Poisson spikes, and that the rest of axons are just the delayed copies of it (not random, since here we assume perfect correlation with no jitter). This reasoning can be repeated for other axons, however, the delayed copies are not going to be just in the future, but will have a set of spikes in the past that will influence the value. Hence, for any axon $a$ the resulting average is simply going to be the sum of $G_{\text{av}}$, plus the responses of the spikes that have already happened, i.e., the expected value for $M_a$ is

$$\langle M_a(x)\rangle_t = C_M \left( G_{\text{av}}(x) + \sum_{t_d^{(i)} > t_d^{(a)}} R(t_d^{(i)} - t_d^{(a)}) \right). \qquad \text{[S14]}$$

This shows that the relative ratios of $\langle M_a \rangle$ are always preserved and dependent only on actual fixed delays, suggesting that the Poisson spiking is more stable and reliable for OMP, since for processes with memory this will not necessarily hold, as is the case for regular spiking, described below, where we show that there is a breaking point when $t_d > \tau_s/2$, since the leading "axon" spikes become the followers.

***Regular spiking.*** In the case of regular spiking, where the ISI times are distributed as $p_{\text{ISI}}(t) = \sum_k \delta(t - \tau_s)$, eq. (6) reduces to evaluating the sum,

$$G_a(\delta t_a) = \sum_{k=0}^{\infty} R(k\tau_s + \delta t_a). \qquad \text{[S15]}$$

The sum in eq. (S15) can be easily evaluated if we note that for our choice of $R(t)$ (eq. (1)), the response function can be written as the difference of two exponential functions, (for clarity, abbreviating $\tau_G$ as just $\tau$).

$$R(t) = e^{-\frac{t}{\tau}} - e^{-\frac{2t}{\tau}}, \qquad \text{[S16]}$$

which reduces to evaluating two geometric sums, yielding,

$$G_A(\delta t|\tau, \tau_s) = \frac{2e^{\frac{\tau_s - 2\delta t}{\tau}} \left(e^{\delta t/\tau} + e^{\frac{\delta t + \tau_s}{\tau}} - e^{\tau_s/\tau}\right)}{\tau\left(e^{\frac{2\tau_s}{\tau}} - 1\right)}, \qquad \text{[S17]}$$

where $\delta t$ is the time to the most recent spike, and $G_a$ is replaced with $G_A$, indicating that the form of $G_A$ will be the same for different axons, but $\delta t$ on separate axons will differ. In other words, the contributions to the global $G(t)$ from different axons is only going to differ through the difference of their times to the most recent spikes, $\delta t_a$, i.e., $G_a(\delta t_a) \equiv G_A(\delta t_a)$. Assuming $t = 0$ coincides with one of the spikes on a given axon we can write, $\delta t \equiv t \pmod{\tau_s}$. When combining $G_A(\delta t_a)$ signals from all axons, generally only one of them can be chosen as such reference, while $\delta t$ for others will be expressed in terms of fixed delays between them, and will require considering separately different orderings, depending on the fixed temporal delays between them. For example, in the simple case, $N_A = 2$, for a given fixed delay, $t_d$, between two regular spiking trains (fig. S1A), we need to distinguish the cases where $\delta t \leq t_b$ *versus* $\delta t > t_b$, where $t_b = \tau_s - t_d$. In figures S1A and B, we color the "leading" axon (the one whose spikes arrive first as "green", while the other one, referred to as "lagging" axon, is colored red. We can see that the time to the last spike will depend on the magnitude of $\delta t_1$. When $\delta t_1 \leq t_b$,

$$G(t) = G_A(\delta t_1|\tau_G, \tau_s) + G_A(\delta t_1 + t_d|\tau_G, \tau_s)$$

and for $\delta t > t_b$,

$$G(t) = G_A(\delta t_1|\tau_G, \tau_s) + G_A(\delta t_1 + t_d - \tau_s|\tau_G, \tau_s).$$

Since $t = 0$ is set for "red" (lagging) spikes, they will occur at times, $t = k\tau_s$, while "green" spikes will occur at $t = k\tau_s + t_d$, and, according to the OMP model, the amount of $M$ will be proportional to $G(t)$ at those times. This is plotted in fig. S1B, and we see that that the lagging axon will receive more myelin promoter than the "green" (leading), which will synchronize them. This, however, switches when $\tau_s < 2t_d$, i.e., for critical spiking frequency of $f_s^{\text{crit}} = 1/(2t_d)$, in this case $f_s^{\text{crit}} = 50$ Hz. Such "reversed myelination" can also be seen in fig. S1B, indicated by "orange" and "red" regions. The dashed lines indicate the contours where the ratios are equal to 0.25, 0.75, and 1, as labeled. The $r_2 = 1$ contour occurs for $f_s^{\text{crit}}$. Evaluating $M_a$ for $N_A$ >2 and in the presence of jitter requires more elaborate calculations, that are left to be addressed outside of the current manuscript.

### Supplementary Figures

While some explanations and the description of the supplementary figures is provided in the main and the SI text, the captions for the SI figures should be detailed enough to be understandable on their own. Figures S1 and S2, supplement the OMP theory sections, both, in SI and in the main text. Figure S4 supports the description of our model selection and fitting (section D). Figures S3, S5 and S6 provide additional information related to the results provided in the main figures (figs. 3 and 4), and figures S7-S10 provide examples of the time progression of OMP variables.

 

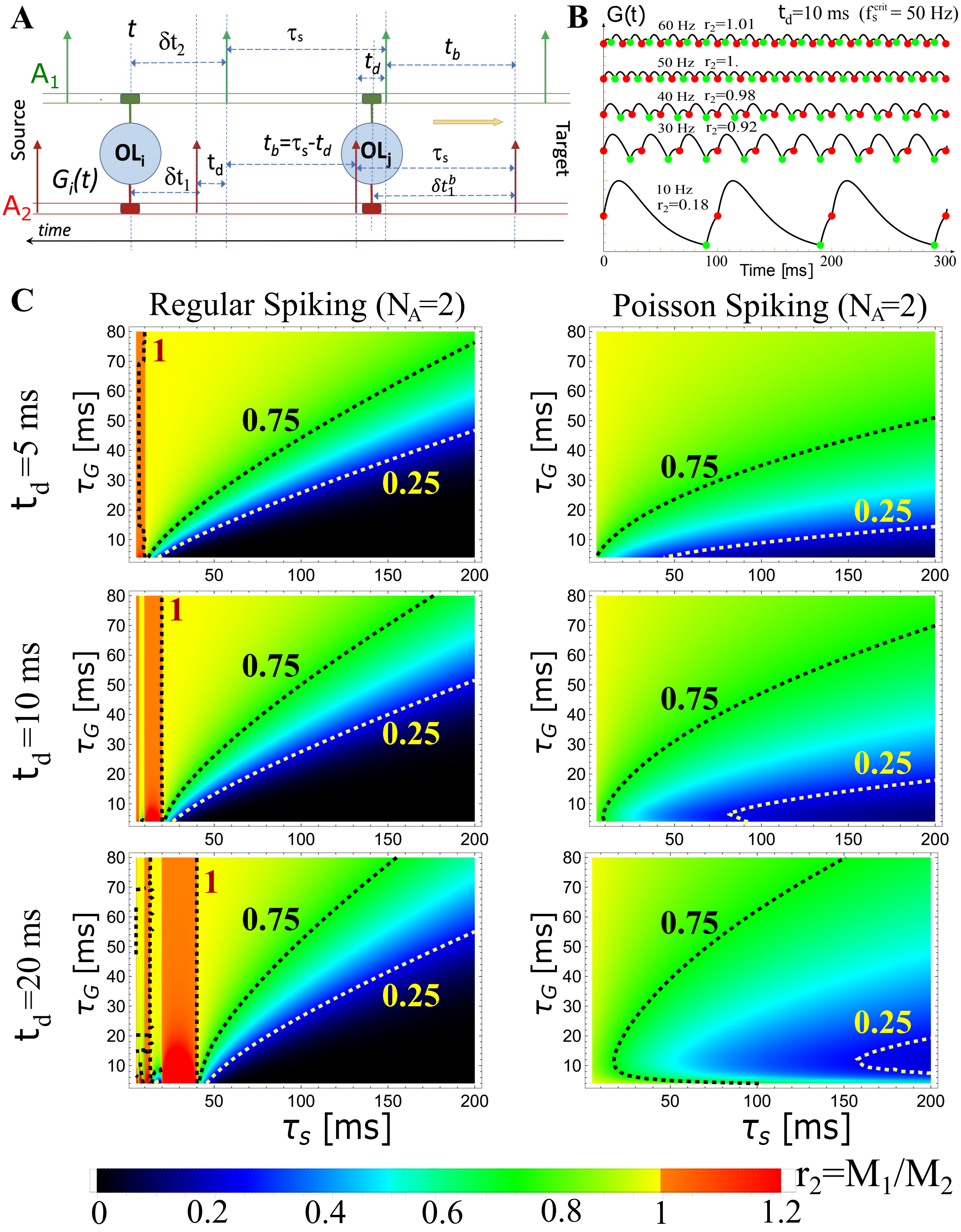


**Fig. S1.** Theoretical predictions for differential concentrations of $M_a$ in different axons, for $\sigma_j = 0, N_A = 2$. (A) Schematic depiction of the temporal relationship between spikes in a pair of axons ($A_1$, $A_2$; $N_A = 2$) in the case of regular spiking, illustrating different scenarios when calculating $G(t)$. The "leading" edge spikes (green) are offset by a fixed delay, $t_d$, over the "lagging" spikes (red). (B) Examples of $G(t)$ for $t_d$=10 ms and for five different spiking frequencies $f_s \in [10, 30, 40, 50, 60]$ Hz, as indicated, showing also the corresponding ratios, $r_2 = M_1/M_2 = M_{\mathrm{green}}/M_{\mathrm{red}}$. Occurrences of "red" and "green" spikes are plotted together with $G(t)$, indicating also the level of $M$ released, as it is proportional to $G(t)$. We see that up to the critical frequency, $f_s^{\mathrm{crit}} = 50$ Hz, the lagging axon will have more myelination factor, $M$, as desired, but this reverses above the critical frequency. (C) Theoretical predictions for the ratio, $r_2 = \langle M_1 \rangle \, / \, \langle M_2 \rangle$, as a function of $\tau_G$ and $\tau_s$ for three different fixed time-delays, $t_d$ =5, 10, and 20 ms (rows, as indicated), in the case of regular (left column) and Poisson (right column) spiking. The regions with red/orange hues are the regions of instability, where $r_2 > 1$, which will lead to further desynchronization, and is only happening for regular spiking. For Poisson spiking, the ratio is always $< 1$, as is evident from eq. (8), hence, the "lagging" axon will always have enhanced myelination relative to the "leading" axon, due to a higher concentration of $M$, which is desired for stable synchronization in all regimes.

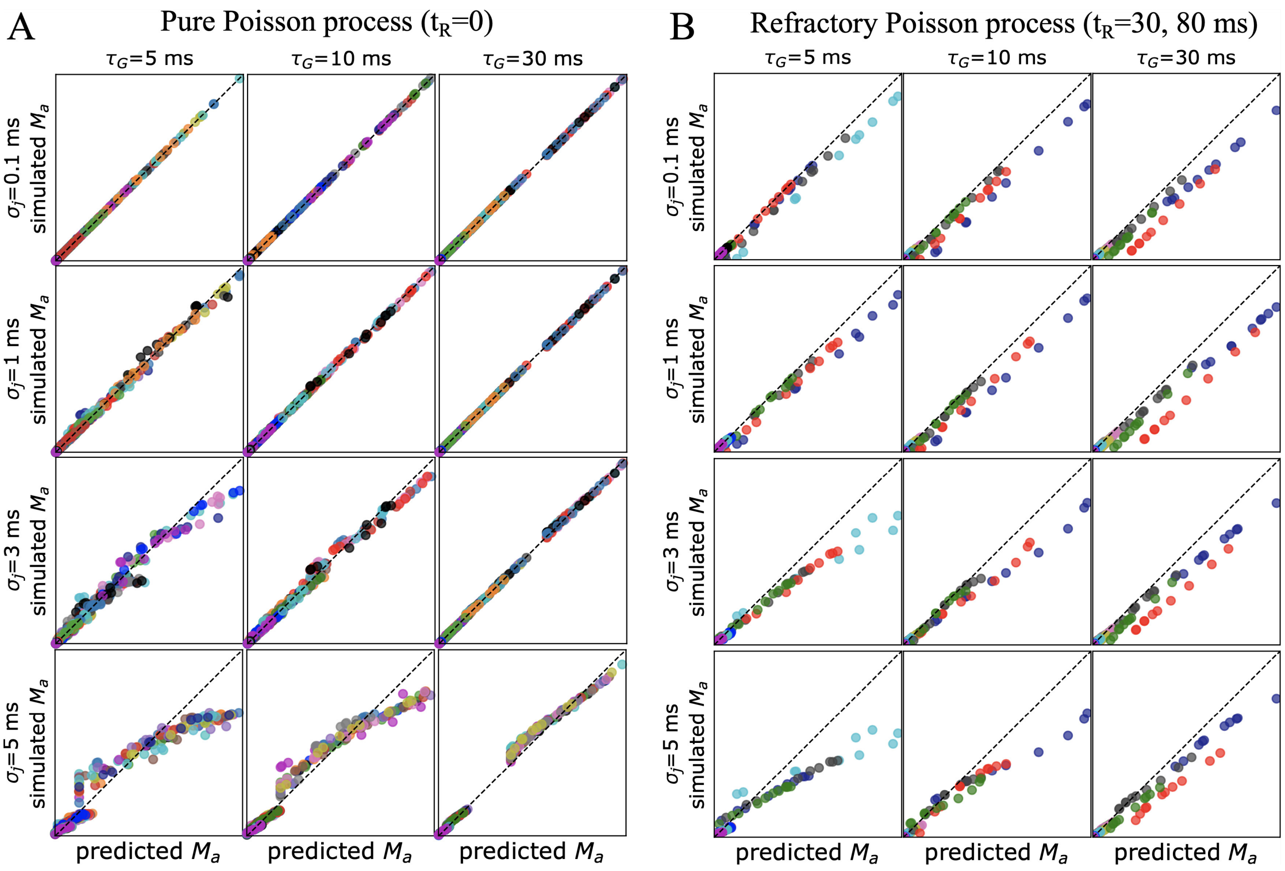


**Fig. S2.** Comparing theoretical predictions with simulations ($\sigma_j > 0, N_A = 10$). The mean $M_a$ values obtained in simulations (y-axis) are compared to the theoretical prediction (x-axis) given by eq. (9), which ignores jitter. (A) The mean $M_a$ values are from a simulation set specified in table S4A, for which correlated spikes with exponential ISI are used (pure Poisson). The results are shown as a grid plot for different values of jitter and characteristic spike response time, $\tau_G$. Individual points from the same parameter set are colored the same, with different parameter sets/runs being colored by cycling through 15 different colors with no particular meaning attached to the coloration. Each set had 2, 5, or 10 points (one per axon). We see that the predictions made for $\sigma_j = 0$ case hold very well for small jitter ($\leq \sigma_j \leq 1$), but fail for larger jitters, particularly for $\sigma_j = 5$ ms, making the differences in $M_a$ across axons smaller. Nevertheless, even for large jitter, when $\tau_G$ is large enough,the prediction given by eq. (9) is in reasonable agreement with the observed/simulated values. (B) Same as in (A), but this time spikes were simulated using a refractory Poisson process with $t_R$=30 and $t_R$=80 ms. Because now it violates the assumption of a memoryless process, under which the derivation of eq. (9) is made, the predictions fail for any amount of jitter or $\tau_G$. We note that theoretical predictions for a general renewal process with any amount jitter are outside the scope of the current manuscript.

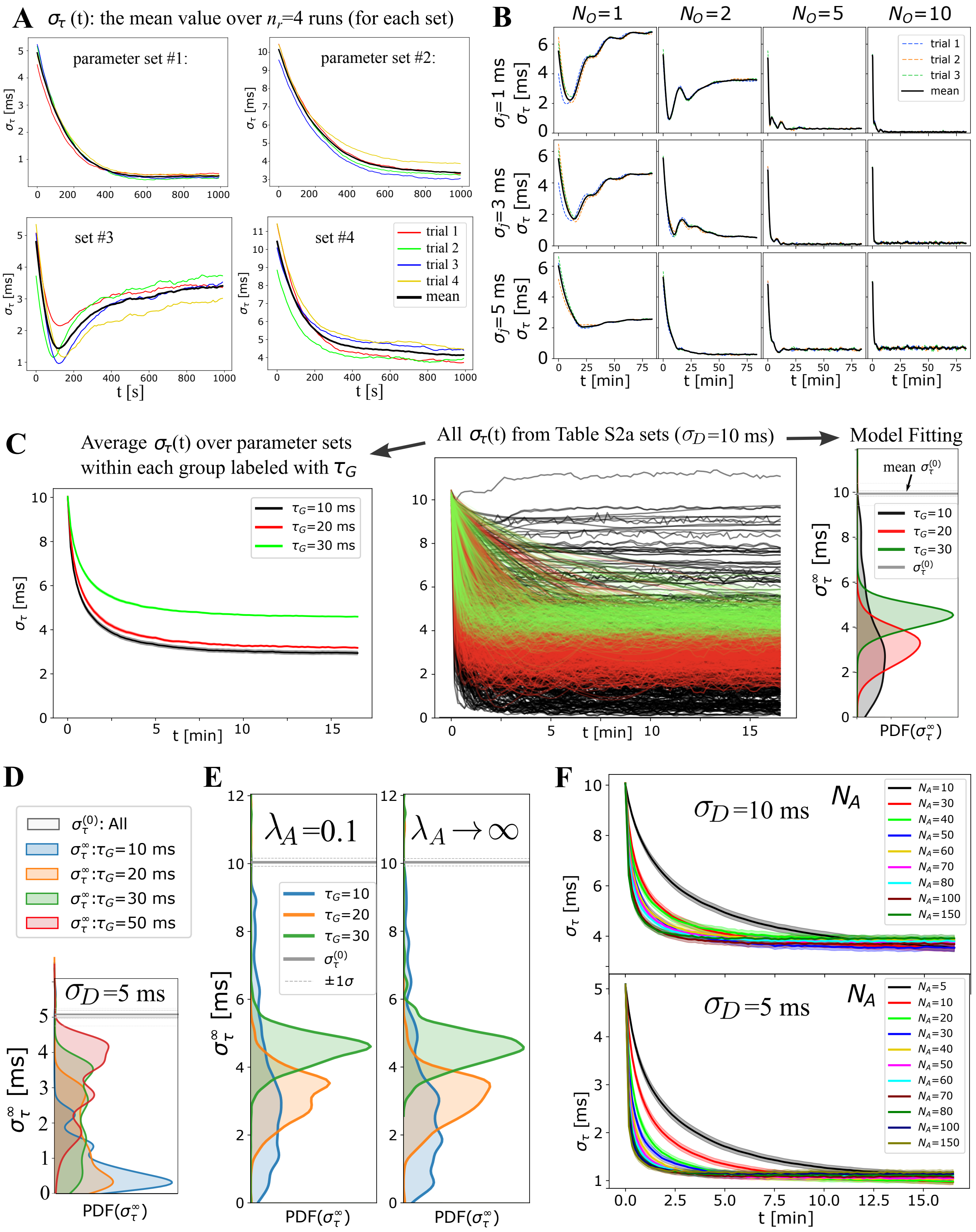


**Fig. S3.** Exploring OMP with a large number of parameter settings (supporting figure for panels A, B, and F in fig. 3). (A) For any given setting of OMP parameters, simulations are repeated $n_r$ times each with random choice of fixed delays ($D_a$, parameterized with $\sigma_D$) and initial adaptive delays (parameterized with $\tau_0$ and $p_\tau$). These individual temporal profiles/trials (colored lines) are averaged and the mean is reported as the synchronization profile, $\sigma_\tau(t)$ (that we refer to as a set or run). (B) A grid plot of $\sigma_\tau(t)$ for correlated spikes using 12 parameter runs (3 trials each), showing its dependence on $N_O$ and $\sigma_j$ (4 $N_O$ and 3 $\sigma_j$ values, for $N_A$ = 10, $\tau_G = 10$ ms, $\lambda_H$=0.000001, $\lambda_M$=0.02). Some of these profiles are shown in fig. 3B to illustrate OMP instabilities with small $N_O$ and fixed delays, which are overcome with increasing $sigma_j$ and $N_O$. (C) The center panel shows all runs specified in table S2a. We summarize such large number of runs in two ways: (1) we average all $\sigma_\tau(t)$ within a subset of runs grouped by some parameter, $\tau_G$, as shown in the left panel; (2) we also fit all $\sigma_\tau(t)$ to 5 models, as described in section D, and plot the distribution of $\sigma_\tau^\infty$ as shown in the right panel. In some cases we also plot the distribution of $\tau_L$ parameters. (D) Main text figure 3A results for $\sigma_D$=10 ms are shown here for $\sigma_D$=5 ms. (E) Main text figure 3A results are compared here to the equivalent runs that were using instantaneous myelination model. We used used a smaller bandwidth for kernel density estimation to highlight the differences, which are not apparent at smoother KDE values. (F) Summary of average profiles grouped by $N_A$, shown in main figure fig. 3F for $\sigma_D$=5 ms, here in addition shown for $\sigma_D$=10 ms.

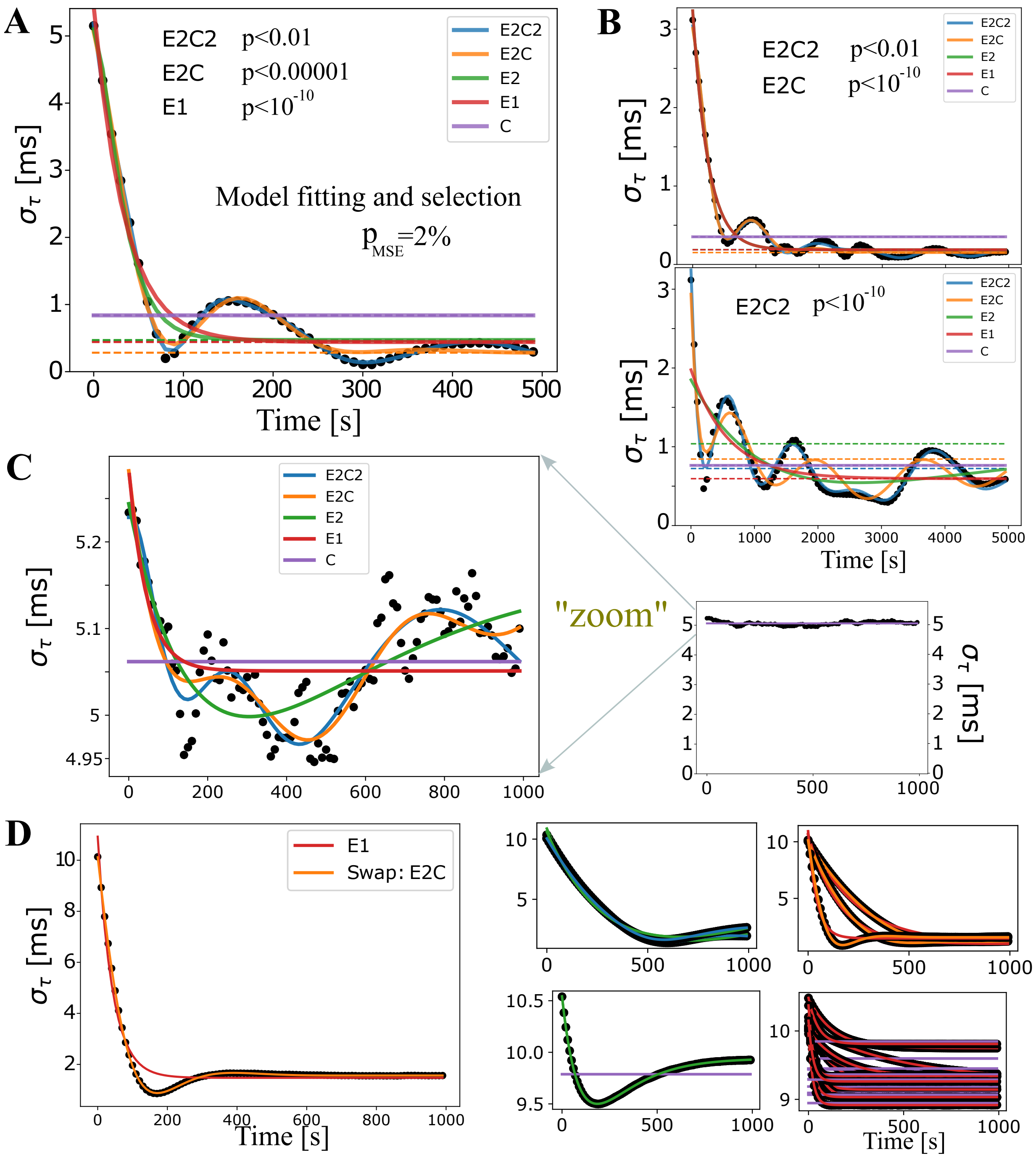


**Fig. S4.** Model fits for quantifying synchronization performance across different parameter runs. (A) An example of fits for all five nested models. Dashed lines represent estimates of the long time baseline, $\sigma_\tau^\infty$. (B) In most cases, comparing the dashed lines, the model selection does not affect the estimate of the baseline (top panel), but can sometimes have a significant impact. For example, in the bottom panel, the baselines differ by nearly 50%. For such a complex synchronization profile it is actually hard to determine the true long time baseline and the only way to resolve those situations is to run really long simulations, which we could not afford in this early and extensive exploratory phase. (C) In the majority of runs, the less restricted models were deemed the best with the standard F-test, and, when magnified, it is clear that E2C2 offers a statistically better description than the constant model, that we label C . On the other hand, when considered at the full scale (small graph on the right), we desire to classify such a profile simply as the constant model, C. This is achieved by introducing a fudge-factor, $p_{\mathrm{MSE}} = 2\%$, which under our modified F-test selects the model C. (D) We also introduced a "swapping" rule. This rule reverts the fit back to the less restricted model when the true MSE in the restricted model is 500 times larger than the one for the restricted model. This did not seem to be consequential, as in most simulations very few swaps occurred, and when they occurred seemed to be justified, as shown in the large panel on the left, where the E1 model suggestion, given by the modified F-test, was reverted to a double exponential fit. The four panels on the right show all swaps obtained with runs specified in table S4c, which was the case which had most the swaps, and most of those were swaps from C to E1 (lower right panel) which had a very small range. Hence, the swap did not change $\sigma_\tau^\infty$ significantly, but had a large effect on the value of $\tau_L$, which is hence less reliable in our summary statistics.

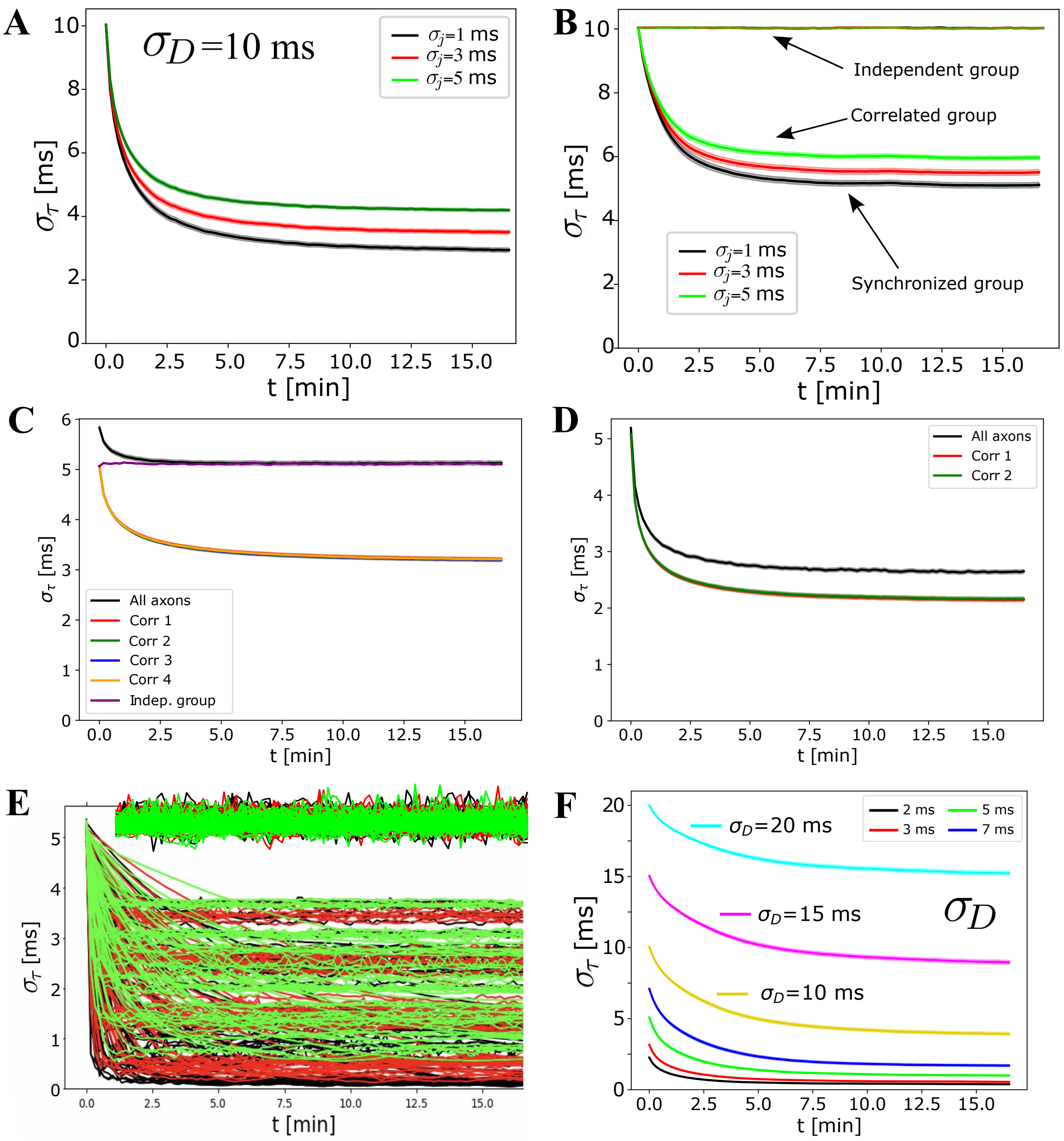


**Fig. S5.** Supplemental result to main text fig. 4. (A-D) Averages over all runs, grouped by jitter, for both $\sigma_D$=5 ms (top row) and $\sigma_D$=10 ms (bottom row), comparing pure signals (left) with mixed signals (right). The shaded region, in the same color, indicates mean $\pm$ SE (standard error) over all runs in a given subgroup (in this case stratified by jitter). (E) Individual runs for the averages reported in (B) where a comparison was made for two groups of axons, one conducting correlated spikes and the other independent spikes. This shows that the actual spread in the performance is much larger than what the shaded standard error (SE) indicates. Averaging over many runs in the case of independent spikes produces extremely small SE (appearing as a "thick" line). Individual runs for independent spikes in (E) are manually offset for clarity as to not overlap with profiles obtained with correlated spikes. (F) Influence of $\sigma_D$ on synchronization efficiency.

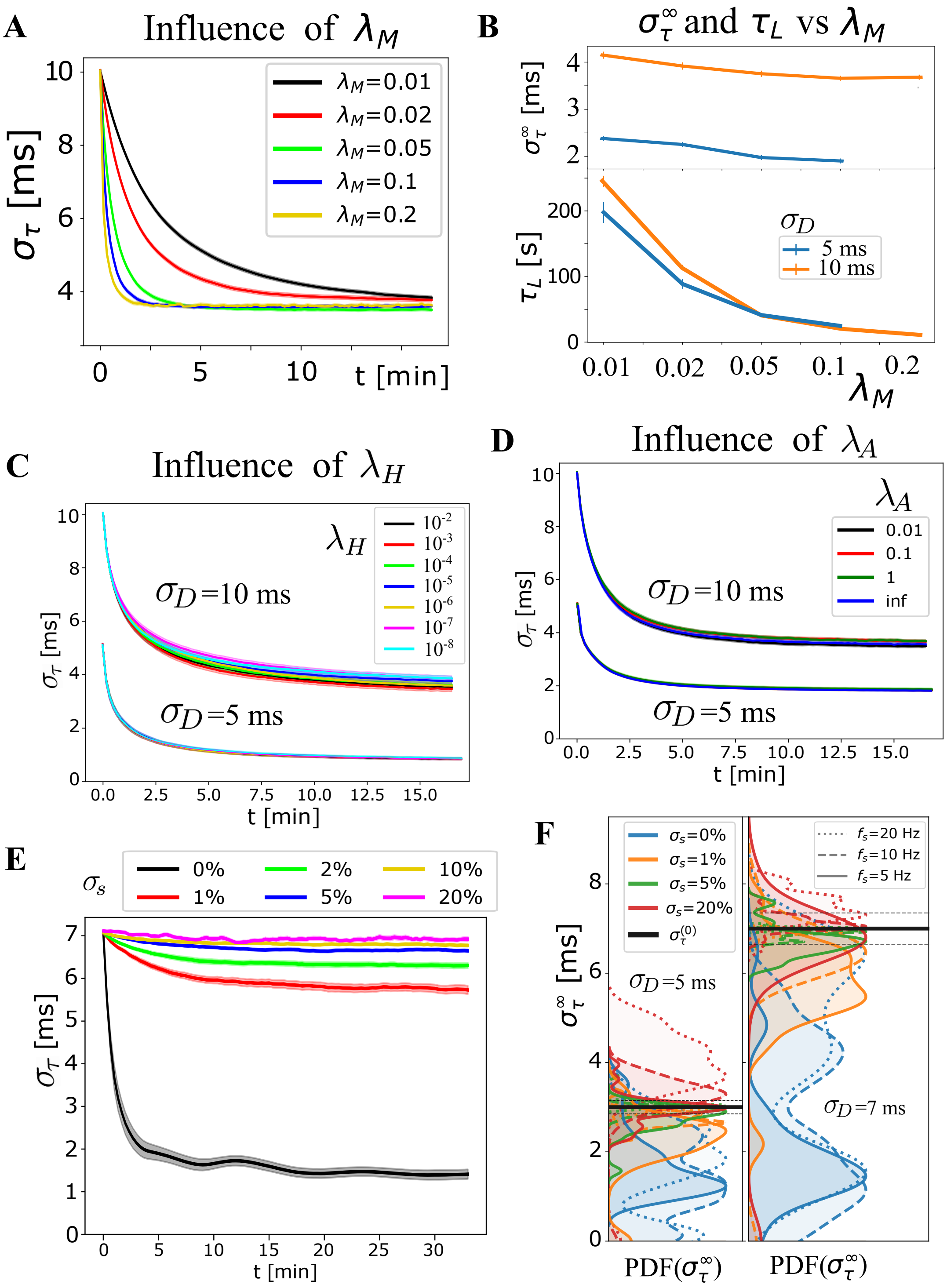


**Fig. S6.** Supplemental figure for exploring the influence of OMP parameters and the rate variability, $\sigma_s$. (A) Average $\sigma_\tau(t)$, grouped by $\lambda_M$, which highly influences the learning rate, as it sets the time scale for changes in myelination. (B) The bottom panel shows this explicitly by using the fitted "learning time", $\tau_L = 1/L_\tau$ (inverse of the learning rate) as a function of $\lambda_M$ for $\sigma_D = 5$ and 10 ms. The dependence of $\sigma_\tau^\infty$ on $\lambda_M$ (top panel) was not strong and it appears to slightly favor faster rates. This "slight" trend does not appear consistently and depends on the range of parameters chosen for averaging. (C) The average $\sigma_\tau(t)$ for $\sigma_D$=10 and $\sigma_D$=5 ms, respectively, show that $\lambda_H$ does not have a strong influence on the behavior of the OMP model (also emphasized in fig. 4E). (D) Average $\sigma_\tau(t)$ for $\sigma_D$=10 and $\sigma_D$=5 ms now grouped by the myelin conversion rate, $\lambda_A$, including the instantaneous myelination model, eq. (13), and showing that $\lambda_A$ has a very small influence on synchronization. This might change when the saturation limits $\tau_{\min}, \tau_{\max}$ become narrower, as well when different $\tau_{\text{nom}}$ values in eq. (15) are used. We allowed the saturation limits to have a broad range from 3 ms to 100 ms, in order to introduce into the model highly heterogeneous values for the delays, making it harder to achieve synchronization. (E) The sensitivity of OMP to firing rate variability is investigated here by introducing a random variation in firing rates, specified via percent deviations, $\sigma_s$. This exploration was limited to regular spiking, as it will produce "beating" synchronized periods, but can also have even longer non-synchronized periods when the OMP dynamics might be detrimental to the arrival time synchrony. The averages of $\sigma_\tau(t)$, grouped by $\sigma_s$, indicate that when the spiking-rate variability is present, but smaller than 5%, synchronization still occurs but becomes very inefficient. This might not be a problem since it is well known that myelin plasticity operates on a very slow time scale, measured in days or even months, compared to synaptic plasticity that sometimes is required to operate on the order of seconds or less. But, it is important that the spread $\sigma_\tau$ does not get worse during non-synchronized epochs, which might not be true for regular spiking. (F) The sensitivity to $\sigma_s$ quantified via the baseline parameter $\sigma_\tau^\infty$. We note that variations greater than 5% can be detrimental, i.e., exhibit de-synchronization effects that are specific to the case of regular spiking and might not arise for Poisson spiking.

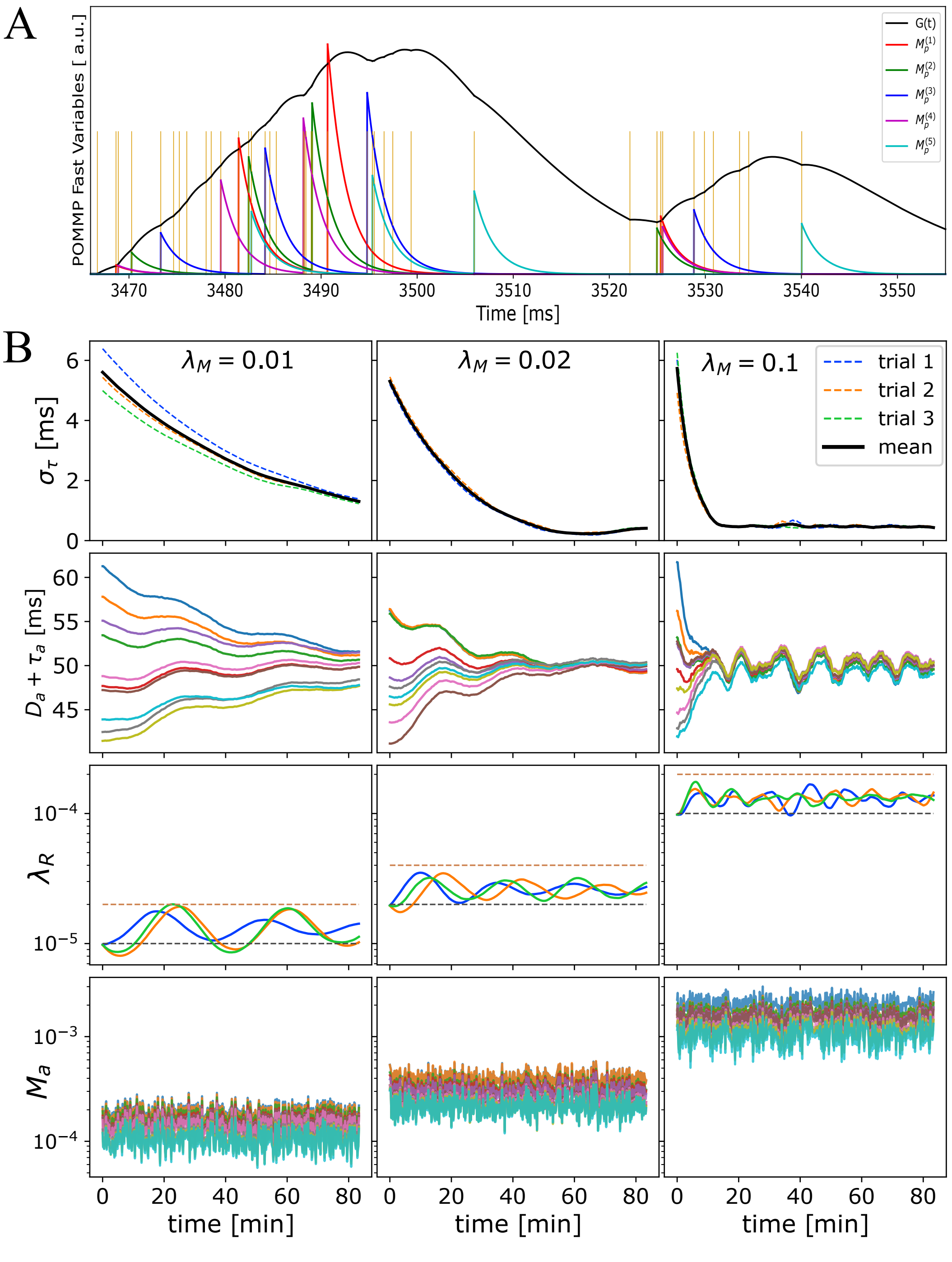


**Fig. S7.** Time progression of OMP variables when correlated spikes with exponentially distributed ISI (Poisson spiking) are conducted through OC with a single OL, $N_O$=1, and for $N_A$ = 10 axons. The ordering of the spikes carries statistically the signature of the fixed delays. The OMP parameters for this simulation are $\tau_G$=30 ms, $\tau_s$=100 ms, $\lambda_H = 10^{-6}$, and $\sigma_j$=5 ms. (A) A millisecond time-scale dynamics of $G(t)$ and $M_a(t)$, with $\lambda_M$=0.01 ms$^{-1}$ and myelination rate $\lambda_A = 0.1$ ms$^{-1}$. For clarity, only 5 of the total $N_A = 10$ axons are shown. Time reported in milliseconds is just an offset from the beginning of the last $T_e$=10 second epoch. (B) Time progression of $\sigma_\tau$ (top row) and OMP model variables, as indicated, over $T_{\text{exp}}$=5000 seconds, plotted as the averages over $T_e$=10 second epochs, and different columns showing different values of $\lambda_M$, as indicated. The second row from top are the 10 delays for each axon, for a single trial, combined with the fixed delays (showing the convergence of the arrival times). The third row shows the time evolution of the myelin removal rate, $\lambda_R$, guided by the homeostatic equation (eq. (15)) with black and red dashed lines indicating the simple homeostatic balance estimate ($\lambda_M N_A/\tau_s^2$) and twice that value, respectively. Differently colored trajectories are the results from three independent trials, with the same parameters but different random initializations of fixed and adaptive delays ($\sigma_{\text{FD}}$=5 ms). The bottom row shows the concentrations of the myelin promoter for each axon in a single trial. For the relatively large, $\tau_G$ = 30 ms, differences in concentrations are not as drastic as they are for $\tau_G$ = 10 ms (see fig. S8).

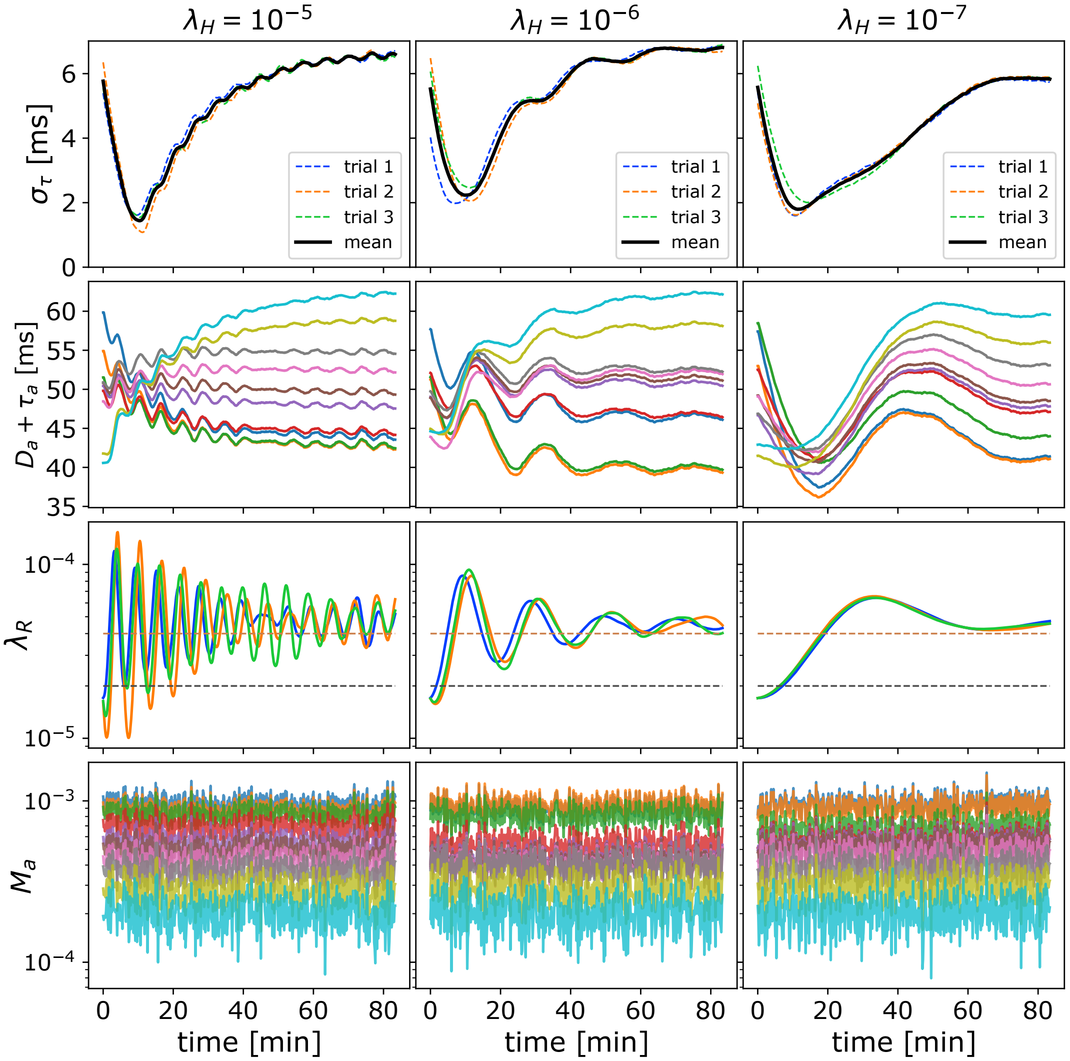


**Fig. S8.** Time progression of the slow variables in the OMP model, similar to one shown in fig. S7B, but now with faster spike-responses in the OL, $\tau_G$=10 ms, and for different $\lambda_H$ (values shown in columns), using only one myelin production rate, $\lambda_M$=0.02, $and\sigma_j$=1 ms. The independent trials, each have with their own set of randomly chosen fixed delays ( $\sigma_{\rm FD}$=5 ms). The top row shows the time progression of $\sigma_\tau$ for each trial (dotted red line is the mean over all trials). The second row are the 10 delays, $\tau_a$, for each axon, and the third row are the same delays when combined with the fixed delays (showing the convergence of the arrival times). The fourth row shows the time evolution of $\lambda_R$, with black and red dashed lines indicating values $\lambda_M N_A/\tau_s^2$ (the balancing estimate for independent spikes) and twice that value, respectively. The bottom row shows the concentrations of the myelin promoter for each axon.

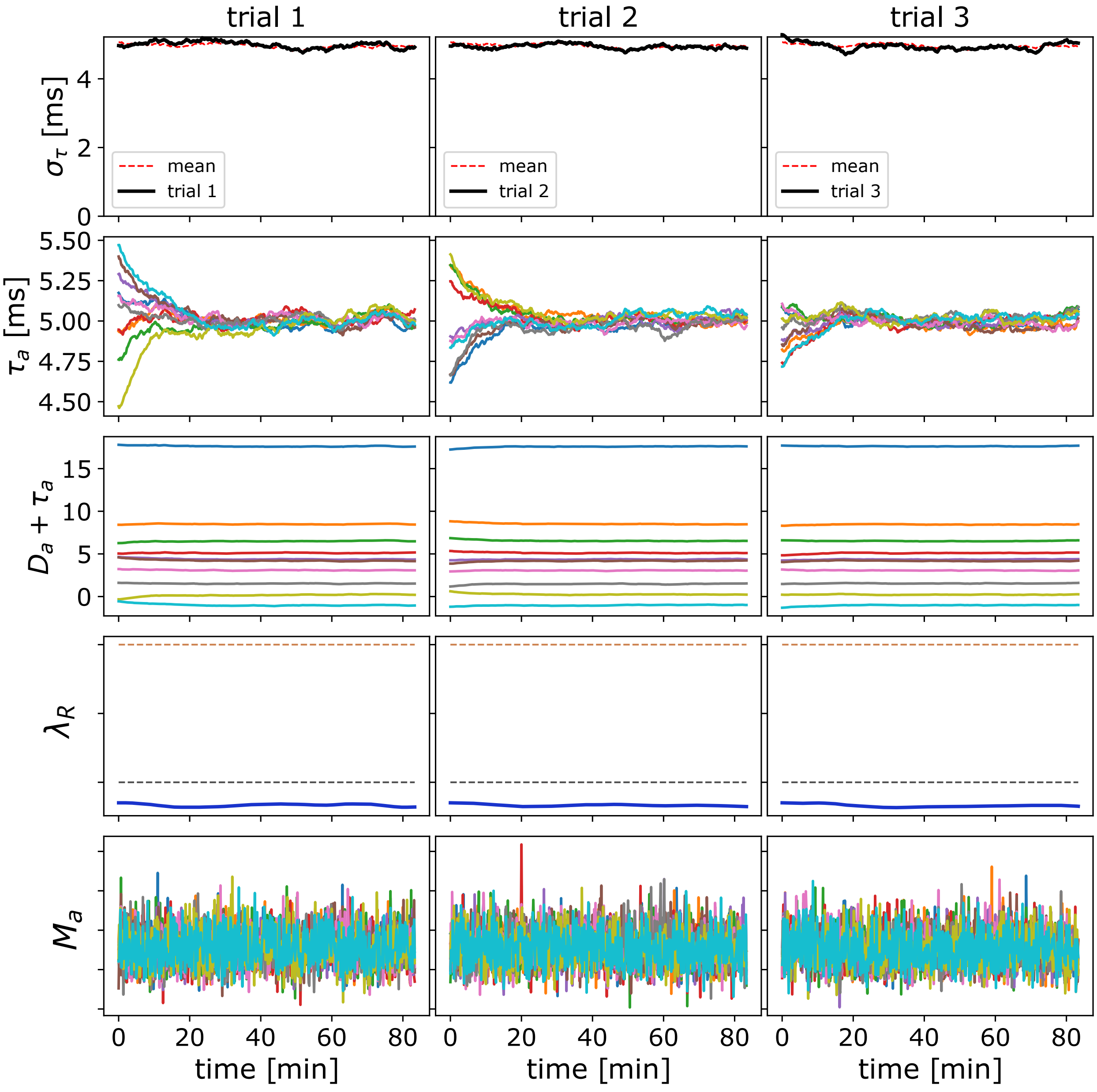


**Fig. S9.** Time progression of the slow variables (averages over $T_e$=10 s epochs) in the OMP model when the spikes are independent with Poisson ISI. Here the columns show three independent trials with the same set of OMP parameters but each with its own set of randomly chosen fixed delays ($\sigma_{\mathrm{FD}}$=5 ms). The OMP parameters for this simulation are $\tau_G$=10 ms, $\tau_s$=100 ms, $\lambda_M$=0.01, $\lambda_H = 10^{-7}$, $\sigma_j$=1 ms, $N_O$=10, and $N_A$=10 axons. The top row shows the time progression of $\sigma_\tau$ for each trial (dotted red line is the mean over all trials). The second row are the 10 axonal delays, $\tau_a$, and the third row are the same delays when combined with the fixed delays (showing the delay profile that will be fed into the next OL segment in the chain)). The fourth row shows the time evolution of $\lambda_R$, with black and red dashed lines indicating values $\lambda_M N_A/\tau_s^2$ (the balancing estimate for independent signals) and twice that value, respectively. The bottom row displays the concentrations of the myelin promoter for each axon.

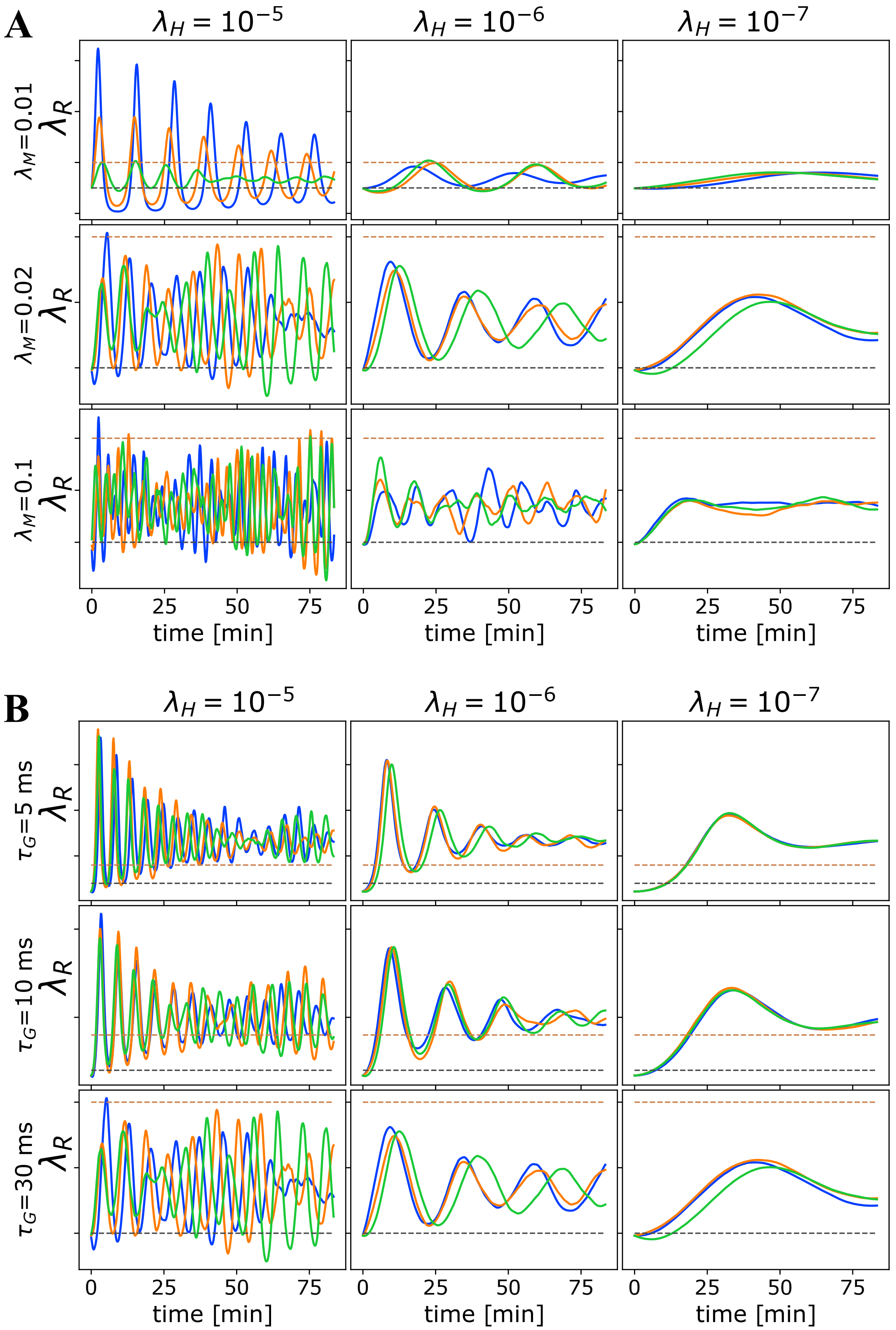


**Fig. S10.** Oscillations in $\lambda_R$ due to homeostatic regulation. The theoretical guess for the balance between addition and removal of myelin, that we initialize $\lambda_R$ to at the start of each simulation, should work well for independent spikes, if saturation effects can be ignored. However, in most cases this prediction does not match the true balancing value for $\lambda_R$, and hence for fast homeostatic rates, the approach to equilibrium will be too fast and will induce oscillations. (A) Time progression of $\lambda_R$ in the OMP model for 3 different values of $\lambda_H$ (columns) and 3 different values of $\lambda_M$ when a set of correlated spikes with exponential ISI (Poisson) are conducted through $N_A$ = 10 axons. For this set of runs $N_O$=1, $\tau_G$=30, and $\sigma_j$=5. (B) Time progression of $\lambda_R$ in the OMP-1 model for 3 different values of $\lambda_H$ (columns) and 3 different values of $\tau_G$ when a set of correlated spikes with exponential ISI (Poisson) are conducted through $N_A$ = 10 axons. For this set of runs $N_O$=1, $\sigma_j$=5, and $\lambda_M$=0.02. The spike-response time, $\tau_G$, does not influence the oscillation frequency of $\lambda_R$ as strongly as $\lambda_M$.

### Tables of Parameters used in Simulations

| Parameter | Values | Unit |
|---|---|---|
| $\lambda_M$ | 0.01, 0.02, 0.05, 0.1, 0.2 | |
| $\lambda_A$ | 0.01, 0.1, 1 | |
| $\lambda_H$ | $10^{-5}$, $10^{-6}$ | |
| $N_A$ | 10, 25 | |
| $N_O$ | 5, 10 | |
| $\tau_G$ | 10, 20, 30 | ms |
| $\tau_{\max}$ | 100 | ms |
| $\tau_{\min}$ | 3 | ms |
| $\tau_{\text{nom}}$ | 50 | ms |
| $\sigma_D$ | 5, 10 | ms |
| $t_R$ | 0 | |
| $\tau_s$ | 25, 50, 100, 200 | ms |
| $\sigma_j$ | 1, 3, 5 | ms |
| $n_r$ | 5 | |
| $n_e$ | 100 | |
| $T_e$ | 10 | s |
| Signals: | Time-Locked & Independent | s |
| # runs | 2× 8640 = 17280 | |

**(a) Parameters for the simulation study exploring the general OMP-1 model behavior for a wide range of its parameters. These same parameters were used for both, the correlated/time-locked spikes and independent spikes. Only simulations for $\sigma_D$=10 ms were used to create Figure 3a; $\sigma_D$=5 ms is shown in figs. S3 and S6**

| Parameter | Values | Unit |
|---|---|---|
| $\lambda_M$ | 0.01, 0.02, 0.05, 0.1 | |
| $\lambda_A$ | 0.01, 0.1, 1 | |
| $\lambda_H$ | $10^{-6}$ | |
| $N_A$ | 10 | |
| $N_O$ | 1, 2, 5, 10 | |
| $\tau_G$ | 10, 20, 30 | ms |
| $\tau_{\max}$ | 100 | ms |
| $\tau_{\min}$ | 3 | ms |
| $\tau_{\text{nom}}$ | 50 | ms |
| $\sigma_D$ | 5 | ms |
| $t_R$ | 0, 30 | |
| $\tau_s$ | 25, 50, 100, 200 | ms |
| $\sigma_j$ | 1, 3 | ms |
| $n_r$ | 4 | |
| $n_e$ | 500/$N_O$ | |
| $T_e$ | 10 | s |
| # runs | 576 | |

**(b) Parameters for the simulations used to create figures fig. 3B,C, and D exploring the influence of the number of OL, $N_O$, in OC.**

| Parameter | Values | Unit |
|---|---|---|
| $\lambda_M$ | 0.02, 0.05 | |
| $\lambda_A$ | 0.01 | |
| $\lambda_H$ | $10^{-6}$ | |
| $N_A$ | 10 | |
| $N_O$ | 5 | |
| $\tau_G$ | 4, 6, 8, ... , 82 | ms |
| $\tau_{\max}$ | 100 | ms |
| $\tau_{\min}$ | 3 | ms |
| $\tau_{\text{nom}}$ | 50 | ms |
| $\sigma_D$ | 10 | ms |
| $t_R$ | 0 | |
| $\tau_s$ | 5, 10, 15, ... 195, 200 | ms |
| $\sigma_j$ | 1 | ms |
| $n_r$ | 10 | |
| $n_e$ | 100 | |
| $T_e$ | 60 | s |
| # runs | 3200 | |

**(c) Parameters for the simulation set used for creating fig. 3E, in which we comprehensively examined the two most important parameters, $\tau_G$ and $\tau_s$.**

| Parameter | Values |
|---|---|
| $\lambda_M$ | 0.01, 0.02, 0.05, 0.1 |
| $\lambda_A$ | 0.01 |
| $\lambda_H$ | 0.000001 |
| $N_A$ | 5, 10, 20, 30, 40, 50, 60, 70, 80, 100, 150 |
| $N_O$ | 5, 10 |
| $\tau_G$ | 10, 20, 30 |
| $\tau_{\max}$ | 100 [ms] |
| $\tau_{\min}$ | 3 [ms] |
| $\tau_{\text{nom}}$ | 50 [ms] |
| $\sigma_D$ | 5, 10 [ms] |
| $t_R$ | 0 |
| $\tau_s$ | 50, 100, 200 [ms] |
| $\sigma_j$ | 1, 3, 5 [ms] |
| nreps | 4 |
| nsreps | 100 |
| Tsecs | 10 |
| Signals | Time-Locked |
| # runs | 4752 |

**(d) Parameters used for OMP-1 simulations when exploring the influence of the number of axons, $N_A$, that OLs myelinate. Runs with $\sigma_D$=5 ms are used in creating Figure 3f. Our OL-axon connectivity was such that $N_A$ was equal to the number of processes each OL extends ($\mathcal{M}$ was a matrix of ones).**

**Table S2. Simulation parameters used in Figure 3; a) used in Figure 3B, C, and D; b) used in Figure 3E; c) used in Figure 3F**

| Parameter | Values | Unit |
|---|---|---|
| $\lambda_M$ | 0.01, 0.02, 0.05, 0.1 | |
| $\lambda_H$ | $10^{-6}$ | |
| $\lambda_A$ | 0.1, 0.01 | |
| $N_A$ | 20, 50 | |
| $N_O$ | 5, 10 | |
| $\tau_G$ | 10, 20, 30 | ms |
| $\tau_{\max}$ | 100 | ms |
| $\tau_{\min}$ | 3 | ms |
| $\tau_{\text{nom}}$ | 50 | ms |
| $\sigma_D$ | 5, 10 | ms |
| $t_R$ | 0 | |
| $\tau_s$ | 25, 50, 100, 200 | ms |
| $\sigma_j$ | 1, 3, 5 | ms |
| $n_r$ | 5 | |
| $n_e$ | 100 | |
| $T_e$ | 10 | s |
| Signals: | TL2, TL+Indep, TL4+Indep | s |
| # runs | 6912 + 2 $\times$ 2304 with pure signals | |

**(a) Parameters for the OMP simulations exploring the situation with mixed signals. The mixed signals were: TL2) - two independently time-locked and synchronized groups, each with $N_A/2$ axons (??C, top panel); TL+Indep) one time-locked/synchronized group and independent, shown in Figure 2a; TL4 + Indep) only 20% axons carry independent and remaining 80% carry 4 equal but separately time-locked/synchronized groups (??C, bottom panel). Two corresponding sets of runs are performed for matched "pure" groups, i.e., purely correlated spikes and independent spikes, matched in terms of $N_A$ involved within groups (for example, the $N_A = 20$ in TL2 will be compared to pure spikes with $N_A = 10$).**

| Parameter | Values | Unit |
|---|---|---|
| $\lambda_M$ | 0.01, 0.02, 0.05, 0.1, 0.2 | |
| $\lambda_A$ | 0.1, 0.01 | |
| $\lambda_H$ | $10^{-2}$, $10^{-3}$, $10^{-4}$, $10^{-5}$, $10^{-6}$, $10^{-7}$, $10^{-8}$ | |
| $N_A$ | 10, 20 | |
| $N_O$ | 5 | |
| $\tau_G$ | 10, 20, 30 | ms |
| $\tau_{\max}$ | 100 | ms |
| $\tau_{\min}$ | 3 | ms |
| $\tau_{\text{nom}}$ | 50 | ms |
| $\sigma_D$ | 10 | ms |
| $t_R$ | 0 | |
| $\tau_s$ | 50, 100, 200 | ms |
| $\sigma_j$ | 1, 3 | ms |
| $n_r$ | 5 | |
| $n_e$ | 100 | |
| $T_e$ | 10 | s |
| # runs | 2520 | |

**(b) Parameters for a set of simulations aiming to explore the influence of homeostatic rate. The results of these simulations are shown in Figure 4D and E and fig. S6**

| Parameter | Values | Unit |
|---|---|---|
| $\lambda_M$ | 0.01, 0.02, 0.05, 0.1 | |
| $\lambda_A$ | 0.1, 0.01 | |
| $\lambda_H$ | $10^{-6}$ | |
| $N_A$ | 10 | |
| $N_O$ | 5 | |
| dmrateinit | 9.e-5 | |
| $\tau_G$ | 10, 20, 30 | ms |
| $\tau_{\max}$ | 100 | ms |
| $\tau_{\min}$ | 3 | ms |
| $\tau_{\text{nom}}$ | 50 | ms |
| $\sigma_D$ | 2, 3, 5, 7, 10, 15, 20 | ms |
| $t_R$ | 0 | |
| $\tau_s$ | 50, 100, 200 | ms |
| $\sigma_j$ | 1, 3, 5 | ms |
| $n_r$ | 5 | |
| $n_e$ | 100 | |
| $T_e$ | 10 | s |
| # runs | 1512 | |

**(c) Parameters for the results shown in Figure 4.e, exploring the influence of $\sigma_D$, i.e., fixed delays on synchronization.**

**Table S3. Simulation parameters used in Figure 4; a) used in Figure 4A, B, and C; b) used in Figure 4D and E; c) used in Figure 4F;**

| Parameter | Values | Unit |
|---|---|---|
| $\lambda_M$ | 0.01, 0.1 | |
| $\lambda_A$ | 0.1, 0.01, 0.001 | |
| $\lambda_H$ | $10^{-7}$ | |
| $N_A$ | 2, 5, 10 | |
| $N_O$ | 1, 5, 10 | |
| $\tau_G$ | 5, 10, 30 | ms |
| $\tau_{\max}$ | 100 | ms |
| $\tau_{\min}$ | 3 | ms |
| $\tau_{\text{nom}}$ | 50 | ms |
| $\sigma_D$ | 5, 10 | ms |
| $t_R$ | 0 | |
| $\tau_s$ | 50, 100, 200 | ms |
| $\sigma_j$ | 0.1, 1, 3, 5 | ms |
| $n_r$ | 3 | |
| $n_e$ | 1000 | |
| $T_e$ | 20 | s |
| # sets | 2916 | |

**(a) Parameters for the simulations used fig. S2A, comparing theoretical predictions for the pure Poisson synchronized spikes with $\sigma_j$=0 to the values obtained in simulations with non-zero $\sigma_j$.**

| Parameter | Values | Unit |
|---|---|---|
| $\lambda_M$ | 0.01, 0.1 | |
| $\lambda_A$ | 0.1, 0.01 | |
| $\lambda_H$ | $10^{-7}$ | |
| $N_A$ | 10 | |
| $N_O$ | 5 | |
| $\tau_G$ | 5, 10, 30 | ms |
| $\tau_{\max}$ | 100 | ms |
| $\tau_{\min}$ | 3 | ms |
| $\tau_{\text{nom}}$ | 50 | ms |
| $\sigma_D$ | 5 | ms |
| $t_R$ | 30, 80 | ms |
| $\tau_s$ | 100, 200 | ms |
| $\sigma_j$ | 0.1, 1, 3, 5 | ms |
| $n_r$ | 2 | |
| $n_e$ | 1000 | |
| $T_e$ | 5 | s |
| # runs | 288 | |

**(b) Parameters for the simulations used fig. S2B, comparing theoretical predictions for the pure Poisson synchronized spikes with $\sigma_j = 0$ to the values obtained in simulations with non-zero $\sigma_j$ and with refractory Poisson spikes, with $t_r = 30$ and 80 ms.**

| Parameter | Values | Unit |
|---|---|---|
| $\lambda_M$ | 0.02, 0.05, 0.1 | |
| $\lambda_A$ | 0.1 | |
| dmrateinit | 9.e-5 | |
| $\lambda_H$ | $10^{-6}$ | |
| $N_A$ | 10, 20 | |
| $N_O$ | 5 | |
| $\tau_G$ | 10, 20, 30, 50 | ms |
| $\tau_{\max}$ | 100 | ms |
| $\tau_{\min}$ | 3 | ms |
| $\tau_{\text{nom}}$ | 50 | ms |
| $\sigma_D$ | 5, 3, 7 | ms |
| $t_R$ | 0 | |
| $\tau_s$ | 50, 100, 200 | ms |
| $\sigma_j$ | 1 | ms |
| $\sigma_s$ | 0, 1, 2, 5, 10, 20 | |
| $n_r$ | 4 | |
| $n_e$ | 100 | |
| $T_e$ | 20 | s |
| # runs | 1296 | |

**(c) Parameters for the simulations used in fig. S6E and F, exploring the influence of the spiking rate variability on the spike synchronization**

**Table S4. Simulation parameter values used for fig. S2 and fig. S6; a) used in fig. S2A; b) used in fig. S2B; c) used in fig. S6E and F.**

### References


1. G Van Rossum, FL Drake Jr, *Python tutorial.* (Centrum voor Wiskunde en Informatica Amsterdam, The Netherlands) Vol. 620, (1995).
2. P Virtanen, et al., SciPy 1.0: Fundamental Algorithms for Scientific Computing in Python. *Nat. Methods* **17**, 261–272 (2020).
3. G Ansmann, Efficiently and easily integrating differential equations with jitcode, jitcdde, and jitcsde. *Chaos: An Interdiscip. J. Nonlinear Sci.* **28**, 043116 (2018).
